# The geometry of the neural state space of decisions

**DOI:** 10.1101/2025.01.24.634806

**Authors:** Mauro M. Monsalve-Mercado, Gabriel M. Stine, Michael N. Shadlen, Kenneth D. Miller

## Abstract

How do populations of neurons collectively encode and process information during cognitive tasks? We analyze high-yield population recordings from the macaque lateral intraparietal area (LIP) during a reaction-time random-dot-motion direction-discrimination task. We find that the trajectories of neural population activity patterns during single decisions lie within a curved two-dimensional manifold. The reaction time of trajectories systematically varies along one dimension, such that slow and fast decisions trace distinct activity patterns. Trajectories transition from a deliberation stage, in which they are noisy and remain similar between the choices, to a commitment stage, in which they are far less noisy and diverge sharply for the different choices. The deliberation phase is pronounced for slower decisions and gradually diminishes as reaction time decreases. A mechanistic circuit model provides an explanation for the observed properties, and suggests the transition between stages represents a transition from more sensory-driven to more circuit-driven dynamics. It yields two striking predictions we verify in the data. First, whether neurons are more choice selective for slow or fast trials varies systematically with the retinotopic location of their response fields. Second, the slower the trial, the more saccades undershoot the choice target. The results highlight the roles of distributed and dynamic activity patterns and intrinsic circuit dynamics in the population implementation of a cognitive task.

## Introduction

Decision-making is a core cognitive process in which the brain integrates noisy sensory information to select an action [Gold and Shadlen, 2007]. Single-cell recordings have been instrumental in elucidating the neural underpinnings of decision-making processes, identifying specific cells that encode abstract rules and decision-related variables [Hanks and Summerfield, 2017, Platt and Glimcher, 1999, Glimcher, 2003, Romo and Salinas, 2003, Carandini and Churchland, 2013].

However, flexible cognitive functions, such as decision-making or spatial navigation, rely on the coordinated and distributed activity of neuronal populations [Vyas et al., 2020, Barack and Krakauer, 2021, Ebitz and Hayden, 2021, Panzeri et al., 2022, Pruszynski and Zylberberg, 2019, Averbeck et al., 2006, Mante et al., 2013, Kiani et al., 2014]. The population dynamics of single trials can be understood as trajectories through a high-dimensional space, the *neural state space*, in which each axis represents the activity of a single neuron in the population. These trajectories are often found to lie, up to noisy fluctuations, on a low-dimensional cognitive manifold within the neural state space [Langdon et al., 2023].

In monkeys trained to communicate their decision with an eye movement, the firing rates of neurons in the lateral intraparietal area (LIP) have been shown to represent the accumulation of evidence for making a saccade toward their spatial response field [Roitman and Shadlen, 2002, Shadlen and Newsome, 2001, Huk and Shadlen, 2005, Shadlen and Kiani, 2013, Churchland et al., 2011, Steinemann et al., 2024]. In the neural state-space framework, these results predict that the manifold associated with each choice should be one-dimensional, such that trajectories on single trials ebb and flow along a line as noisy evidence is integrated. Moreover, averages across trials with different reaction times should produce identical trajectories that progress at different speeds. However, the majority of these previous studies recorded from one neuron at a time and placed the saccade target within the center of the neuron’s response field. As such, it is unclear how interactions across the larger population of LIP neurons, whose response fields span the entire visual field, influence the representation of the decision process.

Here, we show that population-level analysis reveals additional structure beyond that captured by single-cell and cell-averaged studies. We analyze population-level recordings from LIP as monkeys perform a random-dot-motion discrimination task [Stine et al., 2023, Steinemann et al., 2024]. By characterizing single-trial and trial-averaged population activity, we capture the decision manifold, in which the neural trajectories lie, in a data-driven manner and free from prior assumptions about its structure. We find that the decision manifold is inherently two-dimensional and exhibits substantial curvature. One dimension represents the progress of the decision process, from motion onset to saccade preparation. Unexpectedly, reaction time varies systematically along the 2nd dimension, meaning that different reaction times traverse systematically different neural trajec-tories – different patterns of neural activity. The finding suggests structure in the population activity that goes beyond that predicted by previous studies.

Furthermore, we identify two distinct cognitive phases embedded within the decision manifold: deliberation and commitment. In the deliberation phase, trajectories associated with the two choices do not strongly separate; the single-trial trajectories are highly curved and “wiggly”, wander across the reaction-time dimension as well as off the manifold, and move slowly. In the commitment phase, trajectories for the two choices strongly separate, and single-trial trajectories proceed at higher speed directly toward the saccade preparation decision endpoint, with their curvature, “wiggliness”, and wandering strongly suppressed. The difference between the two phases is more pronounced in slower trials, and gradually diminishes as reaction times decrease.

We develop a mechanistic circuit model of the LIP network that accurately reproduces the structure of the decision manifold. The model assumes a retinotopic map across LIP cells. Two key mechanistic insights emerge from the model. First, during deliberation, competing activity bumps representing the two choice targets are retinotopically pulled toward one another by the circuit dynamics. This systematic shift in the neurons involved in the decision, which is greater for longer deliberations and thus longer reaction times, explains the systematic change in neural trajectories with reaction time. Second, the transition to commitment occurs when one activity bump “wins” and grows largely driven by its own self-excitation, representing a transition from sensory-driven to intrinsic-circuit-driven dynamics.

The model makes two key predictions, both of which are validated by the data. First, it predicts that the degree of choice selectivity for single neurons varies from fast to slow trials, with neurons with response fields retinotopically between or outside the targets showing more selectivity for slow or fast trials, respectively. Second, the model provides a behavioral prediction: the slower the trial, the more the monkey’s saccade undershoots the choice target, because the target is represented by the retinotopically displaced activity bump.

Our findings revisit the conceptual framework of neural evidence integration, by illustrating how an interacting and shifting population of neurons integrates evidence in a distributed fashion. The population dynamically and smoothly transitions from a deliberation to a commitment stage as it transitions from a more evidence-driven to more circuit-driven dynamics. This perspective highlights the distributed and dynamic nature of population-level circuit mechanisms underlying decision-making.

## Results

We examined data from large-scale neuropixels recordings from macaque area LIP [Stine et al., 2023, Steinemann et al., 2024]. This study engaged monkeys in a reaction-time motion-direction discrimination task using random dot stimuli. Initially, a fixation dot was centrally displayed, succeeded by two diametrically opposed choice targets. Following this, a random-dot-motion stimulus was shown at the center of the screen (Fig. 1). The direction of motion was always toward one of the two choice targets, and its strength was determined by the motion coherence (see Methods). After an unconstrained deliberation period, termed the trial reaction time, the monkey signified its decision through a saccade to the target corresponding to its perceived direction of dot motion. Reaction times are broadly distributed for any given coherence, but tend to decrease for increasing coherence (Figure 1B), and the percentage of errors decreases systematically toward 0 with increasing coherence (Figure 1C). We remove trials with very long reaction times (top 1%) in all analyses to avoid outliers. In the main text, unless otherwise noted, we analyze one example session with 138 neurons, out of 8 recording sessions in two monkeys; Supp. Fig. 5 shows results from other sessions.

**Figure 1:**
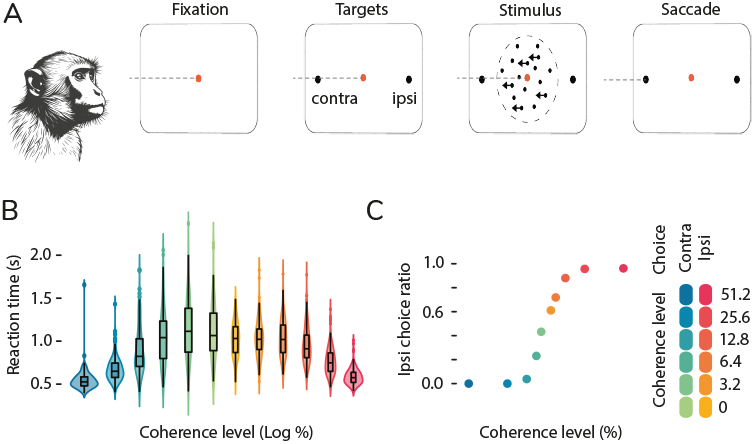
Overview of the Reaction-Time Dot-Motion Direction-Discrimination Task. (A) The task has four phases. In the initial phase, the primate fixates on a central dot, maintaining this fixation throughout the next two phases. During the second phase, two diametrically opposed targets are simultaneously displayed. In the third phase, a circle filled with moving dots is introduced. The dots move in random directions, except at any given time a fixed percentage of the dots move in a common direction toward one of the displayed targets. This percentage, with sign corresponding to contra or ipsi motion, is called the coherence. The monkey studies the motion until at some point it executes a saccade towards the perceived target of the coherent dot motion, with a reward for correct choices (or for a random 50% of trials for 0% coherence). (B)Distribution of the monkey’s reaction times across trials. (C) Fraction of trials in which the monkey selected the ipsilateral (rightward) target, indicating the error rates contingent on coherence.

### The geometry of the neural state space of decisions

Characterizing the geometry of the state space of decisions requires smooth representations of the neural spiking activity at the single trial level. We use AutoLFADS [Keshtkaran et al., 2022, Pandarinath et al., 2018], a deep learning technique, to recover the best guess for the underlying firing rate of a population of simultaneously recorded neurons on single trials (Methods). To correct for representational drift, single trials were aligned in state space at the start of dot-motion stimulation. All analyses are computed in the space of the first ten principal components of the smoothed data (except for the single neuron analysis in Figure 5), but in figures are illustrated in the space of the first three principal components or in a local-geometry-preserving two-dimensional projection described below.

Using a low-pass filter rather than AutoLFADS to smooth the spiking data, we recover our main trial-averaged findings (Supp. Fig. 1). An alternative analysis using a time-warping procedure [Williams et al., 2020] (Supp. Fig. 2) again shows similar trial-averaged behaviors to those we will show with AutoLFADS, but does not permit single-trial analyses.

### The decision manifold is two-dimensional and organized by reaction time

For each choice, single-trial trajectories predominantly reside within an approximately two-dimensional manifold in state space. More precisely, the *intrinsic dimension* of the manifold – the dimension of a local manifold patch – is approximately 2. The manifold is curved, so that its embedding dimension – the dimension of the space containing the full manifold – is higher.

Given the noisy samples, estimating these dimensions is challenging. We estimate the embedding dimension as the participation ratio [Cunningham and Yu, 2014], which has a mean of 2.8 across 8 different sessions for the contra choice (one session illustrated in Fig. 2B) and 3.0 for the complete manifold with two choices. As an alternative to directly measuring the intrinsic dimension, we perform an isomap embedding of the manifold [Tenenbaum et al., 2000]. This maps higher-dimensional data into a lower dimensional space that optimally preserves all pairwise distances, computed as minimal-length trajectories within the data manifold. Examining the reconstruction error of isomap embeddings into spaces of varying dimensions, we find a sharp drop from 1 to 2 dimensions, some improvement in going to 3 dimensions, and very little improvement thereafter (Fig.2B), suggesting that the local geometric structure of these trajectories can be effectively studied within a two-dimensional framework. On this decision-related surface, single trials appear systematically arranged based on their reaction times (Fig.2A), as we will more precisely demonstrate.

**Figure 2:**
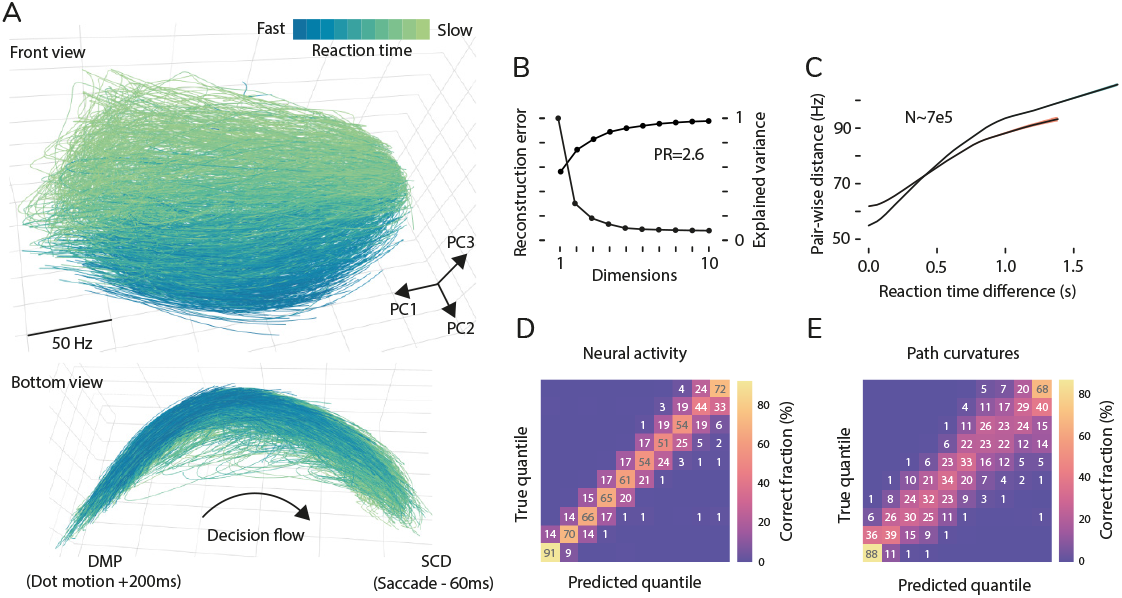
Two-dimensional state space structured by reaction time. (A) PCA visualization of all contralateral trials within state space. Trials are shown for an arc length begining 200 ms after dot motion onset (DMP) and ending 60 ms before the saccade (SCD). Colors indicate reaction time. (B) Explained variance from PCA and reconstruction error from isomap embeddings, plotted against the number of dimensions considered. (C) Distance between pairs of trials in state space as a function of the difference in their reaction times. Lines represent local non-parametric regression for contralateral and ipsilateral submanifolds, with shaded areas (blue, contra; red, ipsi) indicating 95% confidence intervals obtained via bootstrapping. (D-E) Proportion of correctly classified trials by predicted reaction time quantile, using a sequence classifier using either state space trajectories (D) or the first three curvature measures (E).

To compare the geometric properties of space curves, a common approach is to reparameterize these curves using their normalized arc length, a method known as natural parameterization. The normalized arc length is simply the length along the curve in units in which the beginning and end of the curve are 0 and 1, respectively. For each curve, we take its beginning to be the onset of dot motion plus 200ms (DMP), which is the approximate time at which evidence integration begins (Supp. Fig. 2), and the end to be 60ms before saccade onset (SCD), which largely avoids oculomotor preparatory activity. Natural parameterization removes clock time, allowing analysis of the decision process purely in terms of the trajectories through state space, independent of dynamics. It also establishes a common coordinate for comparison of nearby trajectories in state space, assuming that the neural state rather than clock time best captures the current stage of the decision-making process.

To more rigorously assess whether trials are systematically arranged on the manifold by their ultimate reaction time, we conduct several analysis. First, we define the distance between two trajectories as the mean Euclidean distance between pairs of corresponding arc length points. We observe a significant monotonic increase in distance between pairs of trials with increasing difference in their reaction times (Fig. 2C). Next, we study whether a trial’s reaction time can be decoded from its state space trajectory alone, without temporal information. We categorize all trials into ten quantiles based on reaction time and train a sequence classifier to predict the reaction time quantile of a single trial from the state space trajectory (Fig. 2D). The classifier achieves an overall accuracy of 63%, with 96% of predictions falling within either the correct quantile or an adjacent one.

Trials with varying reaction times are distinguished not merely by their locations in state space but fundamentally by their geometrical shape. The fundamental theorem of space curves states that a curve embedded in a high-dimensional space can be fully characterized by its higher-order curvatures, up to Euclidean (shape-preserving) transformations [Carmo, 2016]. Leveraging this theorem, we train a sequence classifier that uses only the first three curvatures of the trial trajectories as a function of arc length (Fig. 2E), without information about the positions of the trajectories in state space. This shape classifier achieves an overall accuracy of 64%, with almost all predictions along the diagonal but with a wider spread compared to the classifier based on the full trajectories, which has access to both shape and positional information. This underscores that trial trajectories with different reaction times are not merely shifted versions of one another but are intrinsically distinct in their geometric properties.

In previous studies, trials have been grouped by motion coherence rather than by reaction time. In Supp. Fig. 7, we show that the manifold shows far less organization by coherence than by reaction time. Reaction time is the result of factors controlling evidence integration on a given trial, such as coherence, but we will illustrate in a mechanistic model how these factors can determine neural trajectories and reaction time by a common mechanism. As such, knowledge of a trial’s ultimate reaction time provides all of the information needed to characterize its expected trajectory.

Is this two-dimensional structure important, or would it be sufficient to analyze the decision process using a locally one-dimensional measure? In the next sections, we show that the two-dimensional structure provides a key framework for understanding integration of evidence in the decision process.

### The decision manifold pattern of deliberation and commitment

Having established that the submanifolds associated with each choice are two-dimensional and organized by reaction time, we now analyze how the overall decision manifold, which encompasses both choices, relates to the decision-making process. To do so, we first determine a trial-averaged trajectory for any given reaction time and choice, using information from the local neighborhood of reaction times. This will allow us to understand how single-trial deviations from the average trajectory influence cognitive processing. To obtain a smooth representation of any scalar variable (e.g., a neuron’s firing rate) vs. reaction time and arc length, we plot the scalar variable from all trials at a given arc length vs. reaction time, and then apply cross-validated nonparametric regression and subsequent interpolation to determine the average value at that arc length for each reaction time (Fig. 3A). Repeating this process for all arc lengths results in a smooth representation of the variable as a function of arc length and reaction time.

**Figure 3:**
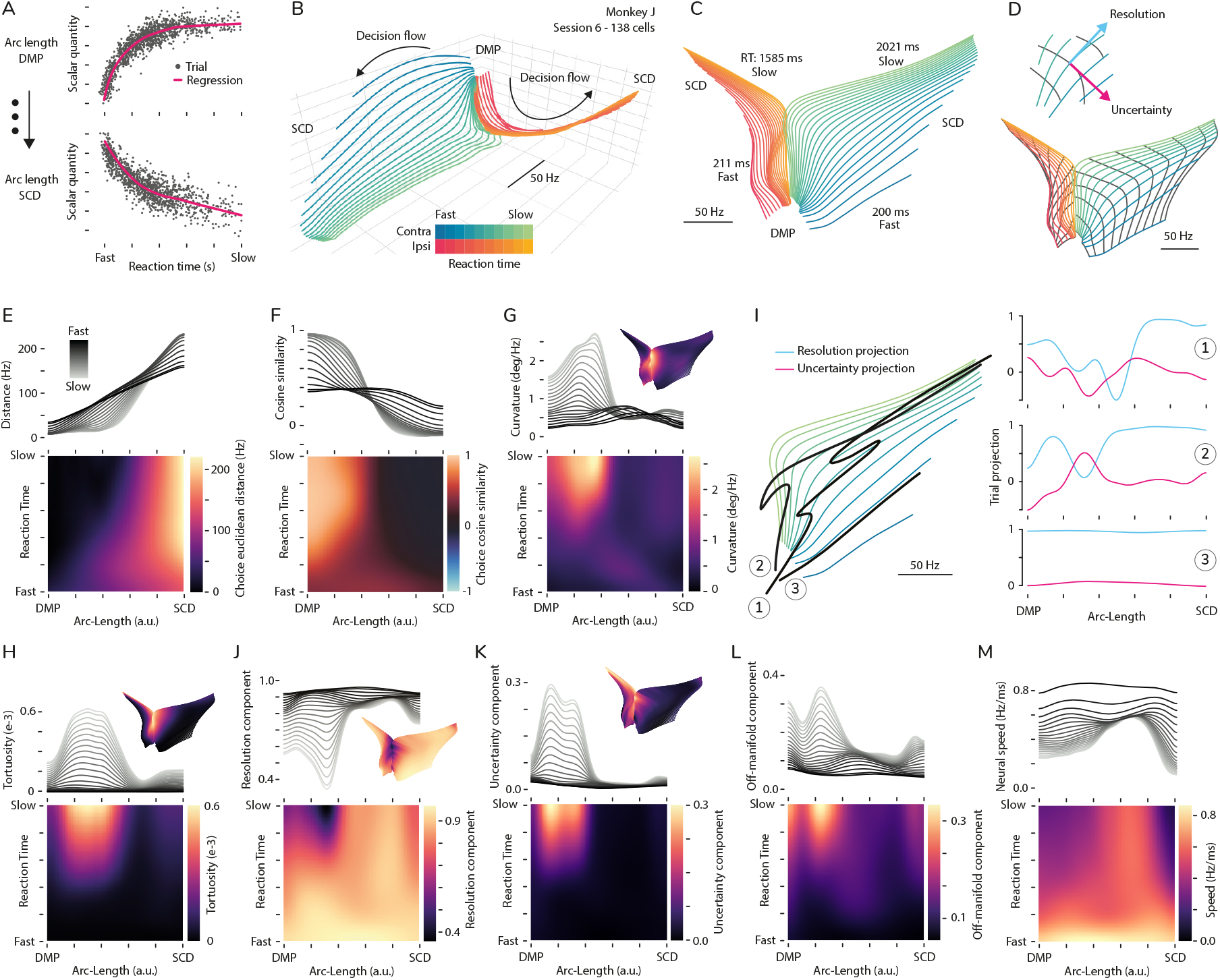
The deliberation-commitment pattern of the decision manifold. (A) Construction of the smooth representation of scalar variables in local embedding coordinates. (B) Linear visualization, in the space of the first three principal components, of the average decision manifold, with lines of constant reaction-time for each of 20 equally spaced reaction times shown in color. (C) Isomap two-dimensional visualization of the average decision manifold. (D) Isomap visualization showing local coordinates (bottom), with lines of constant arc-length in gray. The close-up at the top illustrates the definitions of the resolution and uncertainty vectors; the resolution vector is the unit tangent vector to the mean trajectory. (E-G) Geometric features of the average decision manifold: (E) Euclidean distance between corresponding arc-length points for each pair of mean paths from the two choices with equal reaction-time. (F) Cosine similarity between resolution vectors of the two mean paths, as in (E). (G) Curvature of mean paths. Here and in the following, analysis is of the contra-choice sub-manifold; ipsi is similar. (H-M) Single-trial analyses (quantities computed from single trials, then averaged over trials locally in arc-length and reaction-time, as in (A)). (H) Tortuosity (relative rate of curvature change along the arc-length; “wiggliness”). (I) Left: Example trajectories from three trials (black) superimposed on the mean trajectories (green). Right: Their corresponding tangent vector projections into the local resolution and uncertainty directions derived from the mean trajectories. (J-L) Squared projection of the local direction of change (the tangent vector to the single-trial trajectory) into the resolution (J) and uncertainty (K) directions, and in the direction orthogonal to both (L). (M) Neural speed; this is the only panel to use information from dynamics rather than simply from state-space trajectories.

Using this method to obtain a smooth representation of each neuron’s activity, we first construct a locally-averaged version of the decision manifold, separately for each of the two choice submanifolds. To visualize these submanifolds, we plot the averaged trajectories for 20 equally spaced reaction times from slowest to fastest (Fig. 3B,C). The construction of the average manifold allows us to introduce a natural and interpretable local 2D coordinate system (Fig. 3D). The first coordinate is the normalized arc length introduced above (arc-length). It maps the sequential evolution of the decision-making process in neural state space from DMP to SCD. The second coordinate, reaction-time, leverages the manifold’s organization by reaction time. For each choice submanifold, this coordinate starts at the 97th percentile reaction time (denoted as slow) and ends at the shortest reaction time (denoted as fast).

Upon comparing the submanifolds corresponding to the two choices, we note a pattern in which the slower mean trajectories for the two choices are proximal in state space, and they progress in a similar local direction, reflecting a period of ambiguity or deliberation. Following this initial deliberation phase, there is a pronounced divergence in which the trajectories for the two choices separate sharply, moving in roughly orthogonal directions. The prominence of the deliberation phase, characterized by low separability between the choices, progressively diminishes when comparing trajectories associated with faster reaction times.

We can quantify this trend by comparing pairs of mean trajectories (one for each choice) of equivalent reaction-time values (from slow to fast), in two ways. First, separability is quantified by measuring the Euclidean distance between corresponding points with the same arc-length for the pairs of mean trajectories (Fig. 3E). Second, alignment is quantified by calculating the cosine of the angle between the instantaneous directions of change (tangent vectors) in state space at the same arc-length for the two mean trajectories (Fig. 3F). As reaction times decrease, these measurements indicate an increase in separability and a decrease in early alignment between the trajectories, suggesting that quicker decisions involve a shorter or less pronounced period of neural deliberation.

The transition from the deliberation phase to commitment is marked by a transition in the instantaneous direction of change of the average manifold. One measure of this process is the first-order curvature of the mean paths along the arc-length (Fig. 3G). The curvature at each arc-length point quantifies the local turning angle of the unit tangent vectors, offering a metric for the rate at which the local direction of movement changes. The curvature mirrors the trends observed for the separability and alignment (Pearson correlation coefficients of -0.5 and 0.7, respectively), and peaks at the transition from deliberation to commitment. Note that the mean trajectories are highly curved throughout the deliberation phase, although this is only slightly visible in the 3-D rendering of the trajectories (Fig. 3B) and essentially invisible in 2-D (Fig. 3C). This shows that the decision manifold cannot be summarized by projection onto any one or two constant directions in state space.

We refer to the qualitative dependence of geometric features on the local coordinates as the deliberation-commitment pattern of the decision manifold. We emphasize the gradual dependence on reaction time of this transition. In the following we focus on how fluctuations in single trial trajectories are related to this pattern.

For single trials, the tortuosity of the paths – defined as the relative rate of curvature change along the arc-length – quantifies the “wiggliness” of the curve and indicates the extent to which an individual path is affected by noisy fluctuations. We calculate the tortuosity for all trials and construct its smooth representation in local coordinates (Fig. 3H). The tortuosity metric shows a pattern similar to the deliberation-commitment pattern, further supporting the characterization of the deliberation phase as a stage where the decision path is substantially influenced by variable, potentially noisy cognitive processes. These noisy fluctuations arise from a combination of irrelevant noise and cognitively relevant noise, *e*.*g*. the integration of noisy samples of evidence. To understand the relationship between these fluctuations and the decision process, we analyze how their strength varies depending on their location and direction within the manifold.

Each point in the average manifold has an associated tangent plane, defined by the span of the two unit tangent vectors that follow the curves of constant local coordinates (Fig. 3D). We refer to the local unit tangent vector along the arc-length coordinate as the local “resolution” direction. Conceptually, these unit vectors move along the mean paths of constant reaction-time, so that instantaneous changes in this direction do not correlate with a trial’s reaction time or choice outcome. Similarly, we define the local “uncertainty” direction as the unit vector in the manifold’s tangent plane orthogonal to the local resolution direction and pointing in the direction of decreasing reaction time. Deviations in this direction change the prediction of the trial’s final reaction time or even its choice outcome, especially for trials that are close to the decision separatrix (the line in the manifold of mean trajectories that separates the two choices).

For each trial, at each arc-length point, we compute the projection of the trial’s unit tangent vector onto the corresponding tangent plane of the average manifold at that point. That is, we project the trial’s local direction of change onto the resolution and uncertainty directions (Fig. 3I). we also compute the off-manifold component, *i*.*e*. the part of the instantaneous tangent vector that is orthogonal to the average manifold’s local tangent plane (Fig. 3L). Conceptually, fluctuations in this direction represent the influence of cognitively irrelevant noise. We compute the smoothed representation (Fig. 3A) of the squared components of these projections (Fig. 3J,K); because we are using a unit tangent vector, the three squared components always sum to 1. For slower trials, all of these single-trial projections show a strong pattern of deliberation and commitment. Within the deliberation phase, slower trials show a reduced component along the resolution direction and increased components along both the uncertainty and off-manifold directions. In the commitment phase, slower trials move primarily in the resolution direction, with little movement in the uncertainty or off-manifold directions. Fast trials, throughout their trajectory, behave like slow trials during the commitment phase, moving primarily in the resolution direction.

The pattern seen in the deliberation phase might be, at least qualitatively, expected from evidence integration: the faster the trial, the larger we might expect the deterministic or “drift” component of the evidence to be relative to the noisy, random evidence component and neural noise. The deterministic component should push the trajectory in the resolution direction, while noise in the sensory input would tend to cause excursions in the uncertainty direction and perhaps off-manifold, and neural noise would cause excursions in random directions, most of which are off-manifold. Thus, the faster the trial, the more its instantaneous tangent vectors should point in the resolution direction relative to the other directions. What is striking is that this pattern ceases during the commitment period, when all trials, regardless of reaction time, point primarily in the resolution direction, with excursions strongly suppressed both in the uncertainty direction and off-manifold. This change in slower trials, despite no changes in evidence and no reason to expect changes in neural noise, suggests transition from a phase primarily driven by integration of noisy evidence, to a phase in which trajectories are strongly driven along the decision path to the decision, either suppressing the influence of evidence and neural noise or simply rendering them very small in comparison. This suggests a shift in the neural processing underlying decision-making, moving from deliberation and evidence accumulation to commitment and execution of the choice.

Our analysis thus far has focused on the state space organization of neural activity, without reference to clock time. In Supp. Fig. 6, we show the results of Fig. 3E-H,J-M in terms of time rather than arc length, showing for example that for slower trials the commitment phase typically starts around 500ms before SCD. Finally, we now examine an aspect of single-trial trajectories that depends on clock time, the neural speed – the rate at which the population firing rate vector changes in neural state space. The speed shows a similar division into deliberation and commitment phases (Fig. 3M; Pearson correlation coefficient of 0.7 with the resolution component and -0.6 with the uncertainty component). During the deliberation phase, the speed is slower (deliberation times are longer) for slower reaction times, accounting for much of the difference in reaction times, while during the commitment phase most reaction times converge on a common high speed that then decreases as the target is approached.

Note that, in principle, neural speed could have any pattern without altering the state space geometry of trajectories. The correlation of speeds with geometric trajectories points to a dynamical mechanism shaping the geometry of decision-making, as we now examine in a mechanistic circuit model.

### A mechanistic cortical model of visual decision-making in LIP

To understand how our previous observations might be mechanistically implemented by neurons in the lateral intraparietal area (LIP), we developed a mathematical model based on excitatory and inhibitory neurons representing retinotopic position in the visual field on a two-dimensional retinotopic sheet, with the strength of connectivity between neurons decreasing with retinotopic distance (Fig. 4A). Note that nothing about choice or the decision process is explicitly built into the circuit. A key feature of the model is the non-linear interaction between neurons, implemented through a rectified power-law input-output relationship; the specific model we present uses power 2 for inhibitory neurons and 1 for excitatory neurons (see Methods).

**Figure 4:**
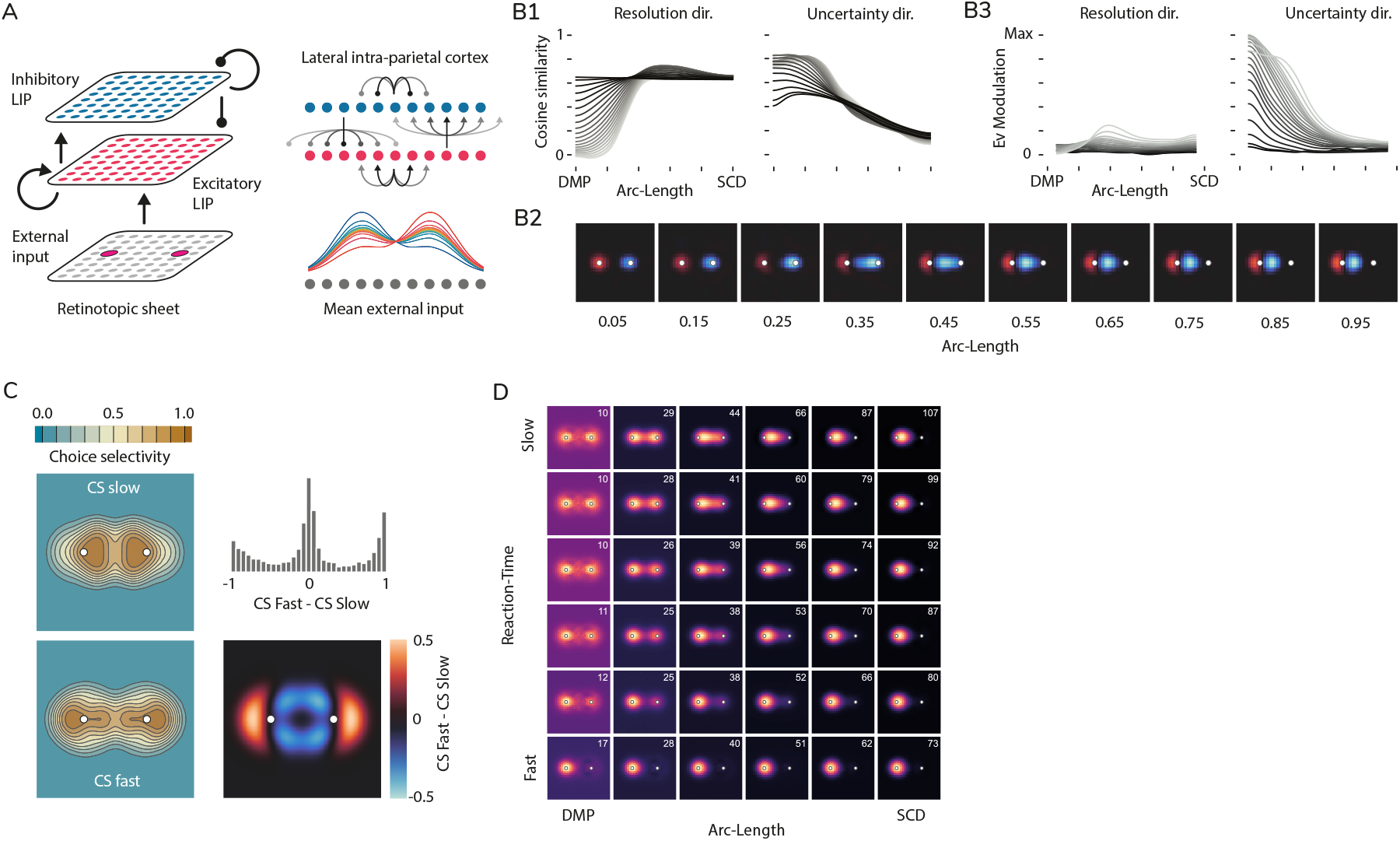
Implementation of the decision-making process in the LIP model. (A) Illustration of the twodimensional retinotopic sheet model representing the lateral intraparietal area (LIP) based on the retinotopic representation of the visual field. The model features excitatory and inhibitory neurons with connectivity strengths that decrease with cortical distance, incorporating non-linear interactions and external inputs that simulate visual and cognitive processing during decision-making tasks. In the following all calculations are done on the excitatory cell population activity. (B) Analysis of evidence integration in the model. (B1) Cosine similarity between the direction of evidence for the contra choice and the local resolution and uncertainty directions in the average contra submanifold. (B2) Local uncertainty-driving input directions for the slowest reaction-time window for varying arc lengths. (B3) Evidence modulation: Projection of the local resolution-driving and uncertainty-driving input directions onto the global evidence direction for the contra choice. (C) Neuronal choice selectivity in the model as a function of retinotopic location (left), illustrating differences in selectivity for slow (cognitively challenging) versus fast paths. The difference map (bottom right) reveals retinotopic clustering of neurons that are more choice-selective neurons for the slowest or fastest mean paths. The upper right histogram displays the distribution of the difference in selectivity between fastest and slowest paths among neurons with a selectivity score of at least 0.5 for both fastest and slowest paths. (D) Firing rate maps on the retinotopic sheet, illustrating the progression of the decision process within the manifold’s local coordinates. Displayed are maps corresponding to six equally spaced points along both arc-length and reaction-time in the manifold. The top right corner of each map indicates the peak firing rate in Hz, with the colormap ranging from 0 Hz to the peak. Because decision threshold is the mean activity over cells within a circle about the target, the peak rate at threshold (rightmost column) is higher with greater displacement of the activity bump from the target.

Neurons receive external inputs that encode information about the strength of evidence for each choice. The mean inputs are modeled as two constant activity bumps centered at the visual field locations of the two targets (Fig. 4A), assuming that top-down attention is focused on these locations, with relative amplitudes linearly modulated by the signed coherence level of the dot motion stimuli. In addition, excitatory cells receive stimulus-independent, spatially correlated pink noise to account for the joint variability arising from noisy evidence and inherent fluctuations in neural activity.

We declare a choice to be made when the absolute difference between the mean activities in circles of cells centered on the two targets reaches a threshold value. The model qualitatively captures the psychophysical curves associated with the task, *i*.*e*. the distributions of reaction times and the error rates as a function of coherence level (Supp. Fig. 3).

We applied the same analyses to the model data as we did to the empirical data (Supp. Fig. 3). The model, like the data, produces a decision manifold that is locally two-dimensional but non-planar, organized by reaction time, and visibly shows the separation of deliberation and commitment phases. The model reproduces all of the features of Fig. 3E-G,H-M that more quantitatively characterize the cognitive pattern of deliberation-commitment, both in the average manifold and in the single-trial analyses (Supp. Fig. 3). Running the model without noise for nonzero coherence reproduces the average manifold (not shown), suggesting that the resolution direction represents the local direction of growth under constant evidence and intrinsic circuit dynamics, while fluctuations in the uncertainty direction are driven by fluctuations in momentary evidence, whose influence is suppressed in the commitment period.

How does the direction of growth of the population pattern of activity relate to the direction driven by the input evidence? To examine this, we define the global evidence direction for the contra choice – the direction driven by the input evidence – as the difference between the trial-averaged input activity vectors for contra and ipsi alternatives for coherence 0. By calculating the cosine similarity between this global direction and the local resolution and uncertainty directions of the average contra submanifold (Fig. 4B1), we find that during deliberation, the initial resolution vectors are orthogonal to this global evidence direction for the slowest mean paths, but are increasingly aligned with the global evidence vector for decreasing reaction-time. During the commitment phase, resolution vectors for all mean paths show high alignment with the global evidence direction, regardless of reaction time. In contrast, the uncertainty vectors display the opposite trend: alignment is high during the deliberation phase, highest for the slowest paths, and gradually decreases and becomes independent of reaction-time in the commitment phase.

However, alignment of the global evidence direction with the local resolution or uncertainty direction does not imply that the global evidence is actually driving the neural trajectories along those directions. Alternatively or additionally, the network’s internal dynamics may be driving the trajectory along a given direction. To disambiguate the relative roles of these two factors, we examine how momentary evidence fluctuations are integrated at the single trial level. We find local resolution-driving and uncertainty-driving input directions, *i*.*e*. local directions in state space whose projections onto the fluctuations of the full input vector from its mean best predict the resolution and uncertainty components, respectively, of the unit tangent vectors of single trials (see Methods). The local uncertainty-driving input direction (Fig. 4B2) reveals a drastic shift in how the model integrates evidence. For slow trials, early in deliberation the local uncertainty-driving direction closely aligns with the global evidence direction, but progressively decreases in alignment during the evolution of the process to become largely orthogonal to this global evidence during commitment.

To characterize this alignment throughout the manifold, we compute an evidence modulation measure as the projection of the local input directions into the global evidence direction (Fig. 4B3). We observe that, despite the high alignment during commitment of the local resolution direction of the average manifold with the global evidence direction (Fig. 4B1), the local resolution-driving input direction shows little alignment with the global evidence direction (Fig. 4B3) for any reaction-time at any decision phase, meaning that momentary evidence does little to drive activity in the resolution direction. On the other hand, the local uncertainty-driving input direction exhibits high alignment with the global evidence early in single trials for slower trials, which progressively diminishes during deliberation until it reaches a stationary level during commitment, similar to the levels observed for resolution. The strength of this initial alignment decreases with decreasing reaction-time in the manifold.

These findings suggest two key points about the model, which serve as predictions: first, momentary evidence is integrated almost exclusively along the uncertainty direction, and second, this integration primarily occurs during the deliberation phase.

We next examine choice selectivity in the model. We define a neuron’s choice selectivity based on the Pearson correlation between its contra and ipsi activities in mean paths with equivalent reaction times during the commitment phase. Choice selectivity is zero when the activities, across arc-length during the commitment phase, are perfectly correlated (*e*.*g*., both are rising or both are falling during the commitment phase), and one when they are perfectly anticorrelated. Comparing choice selectivity as a function of a neuron’s retinotopic position for the slowest and fastest paths, we see that a neuron’s choice selectivity depends both on the trial’s ultimate reaction-time and the neuron’s retinotopic position (Fig. 4C, left). In particular, the map of differences between choice selectivities in fast and slow paths (Fig. 4C, bottom right) shows a striking pattern: neurons with retinotopic position near the choice targets exhibit no difference in selectivity between the reaction times, neurons with positions displaced from a given target towards the middle show more selectivity for slow paths and neurons displaced away from the middle show more selectivity for fast paths. This suggests that in previous single-cell experiments, in which the target is placed within the center of a cell’s response field, changes in choice selectivity based on reaction time would not be observed. Another way to view these results is to look at the distribution across selective neurons (*CS >* 0.5) of the difference in their choice selectivity for fast and slow paths (Fig. 4C, upper right). Selective neurons cluster at differences of -1, 0, and 1, reflecting distinct subpopulations of neurons that either show strong selectivity for one reaction time while being weakly selective for the other (-1 or 1) or maintain consistent selectivity across both (0).

The origin of these model behaviors is in the evolution of the activity in the retinotopic sheet as the input is integrated. Consider trials that end in a contralateral choice (Fig. 4D). The input drives initial activity centered on each target. On the fastest trials, the bump of activity centered on the contra target quickly suppresses the ipsi-target-centered activity and rises quickly to threshold, strongly driven by its own self-excitation, without moving its location. Already at DMP the trial appears to be in the commitment phase, with trajectories rapidly proceeding along the decision path. On longer trials, the two bumps are more equal and compete, producing the deliberation phase. Each bump’s activity excites cells nearer to it and suppresses cells further away, pulling each bump towards the other. The longer the deliberation phase persists, the further the bumps move. The commitment phase arises when the contra bump gains the advantage, suppressing the ipsi bump while rising rapidly at its displaced location to a decision. This explains why the cells displaced toward the middle from the contra target are primarily choice selective on slower trials, and those on the opposite side on faster trials. The 2-D decision manifold, its non-planar curvature, and the various measures of Fig. 3 follow in the model (Supp. Fig. 3) from the noisy evolution of the moving activity bumps.

### Testing model predictions: Neural and behavioral signatures of the competitive dynamics

The gradual shift in activity-bump location with increasing deliberation time leads to two key predictions. First, there should be a relationship between a neuron’s choice selectivity, the location of its response field, and the trial’s reaction time. To test this prediction, we first compute the choice selectivity of neurons separately for the slowest and fastest reaction times. Our analysis reveals that, for either group of reaction-time, neurons clustered into high and low selectivity groups. Furthermore, subgroups of neurons shifted their selectivity from high to low or vice versa depending on reaction-time(Fig. 5A,B,C). To examine dependence on response field locations, for the fastest and the slowest mean path, we construct a visuotopic mapping of choice selectivity as a weighted average of the neurons’ response fields, weighted by their choice selectivity. These mappings exhibited significant visuotopic clustering of choice selectivity for both groups of reaction-times (Moran’s *I >* 0, permutation test, *p <* 0.001). The maps were also significantly different from each other (2D Wasserstein distance, permutation test, *p <* 0.001), indicating a shift in the recruitment of neurons for slow or fast paths depending on the locations of their response fields, as predicted by the model. Examining the difference between fast and slow visuotopic maps indicates that choice-selective cells tend to cluster toward the center of the visual field for slow paths, and away from the targets for fast paths (Fig. 5C), as the model predicts(Fig. 4C).

**Figure 5:**
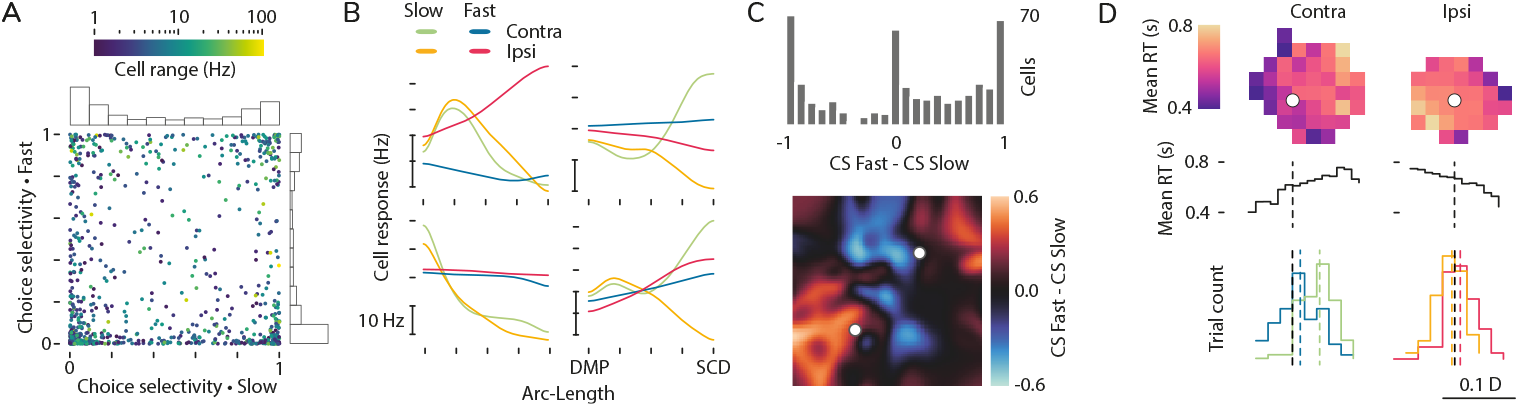
Tests of model predictions. (A) Choice selectivity computed for the slowest and fastest mean paths for all cells across all 8 sessions with a dynamic range in any of the 4 mean paths (2 reaction times, 2 choices) of at least 1 Hz (N=723). (B) Activity of four example cells from each corner of the plot in (A), with panels positioned accordingly. Each panel shows the activity of the cell for both choices and both the slow and fast paths. Vertical scale bars indicate 10 Hz. (C) Histogram showing selectivity changes for neurons with at least a 1 Hz range and restricted to a choice selectivity of 0.5 or higher in either the slow or fast paths across all sessions (top, N=484). The lower panel displays the difference for the example session between the visuotopic maps of choice selectivity for fast and slow paths (bottom). (D) Relationship between saccade end-point locations and reaction time across trials for the example session. Contra and ipsi choices are shown side by side for comparison. The bar at the bottom represents the scale relative to the distance *D* between the targets. Target locations in retinotopic space (white circles) were determined from the visually guided task by averaging eye positions across trials within a 10 ms window immediately following saccade completion. Top: Mean reaction time in each bin of saccade end-points for each choice. Middle: Mean reaction times for projected locations onto the line connecting the targets. Black dashed vertical lines indicate projected target locations. Bottom: Distributions of projected locations for trials in the top and bottom deciles of reaction time. Colors as in (B), *i*.*e*. darker is fast, lighter is slow. Colored dashed vertical lines indicate the medians.

The model makes a second, behavioral prediction that we can test with the existing data. If we assume that the location of the activity peak in LIP influences the saccade endpoint, as suggested by much previous work [*e*.*g*., Bisley and Goldberg, 2010], then the gradual displacement of activity toward neurons with response fields that are between the targets should give rise to saccades that undershoot the target on trials with slower reaction times. To test this, we computed the saccade endpoint for each trial in the example session and computed the average reaction time for each small bin of endpoints. The result, as predicted, is a systematic shift from slower to faster reaction times with increasing eccentricity of the saccade endpoint (Fig. 5D, top; spatial organization of reaction times is significant, 2D Wasserstein distance, permutation test, *p <* 0.001 for both choices). To more quantitatively assess the directionality of this organization, we projected the 2D saccade locations onto the line connecting the two targets and computed the average reaction time in each 1D bin. This showed a systematic decrease in reaction time with increasing eccentricity (Fig. 5D, middle; Spearman correlations between reaction time and projected location, 0.9 and -0.8 for contralateral and ipsilateral choices, respectively). Alternatively, we can measure this effect by comparing the distributions of projected saccade locations for the top and bottom deciles of reaction times for each choice (Fig. 5D, bottom). The distributions are significantly different (Kolmogorov-Smirnov two-sample test, *p <* 0.001 for both choices), with the median of the slowest decile shifted toward the opposite target. As a control, the distributions of saccade locations projected along the axis orthogonal to the inter-target line show no significant differences between the top and bottom reaction time deciles. For other sessions, we lacked data sufficient to reconstruct target locations, but for each session and choice, we compared the 2D distributions of saccade locations between the fastest (bottom decile) and slowest (top decile) reaction time trials. Using a permutation test based on the 2D Wasserstein distance, we observed significant differences *p <* 0.05 in 12 out of 16 datasets (2 choices across 8 sessions).

## Discussion

Our study provides new insights into the population dynamics of decision-making in the lateral intraparietal area (LIP), expanding upon a conceptual framework in which single neurons are thought to directly represent the accumulated evidence for making a saccade toward their response field. In previous work, neurons were typically recorded one at a time with the choice target within the center of a neuron’s response field. This paradigm generated numerous insights into how the decision process was represented in LIP [Gold and Shadlen, 2007, Shadlen and Kiani, 2013]. However, a larger-scale perspective has been lacking on the role of the full population of LIP neurons in the decision process, particularly given their broad and heterogeneous response fields [Blatt et al., 1990, Barash et al., 1991]. By analyzing population recordings of multiple LIP neurons with the target fixed at a single location [Stine et al., 2023, Steinemann et al., 2024], we study neural activity spanning the entire visual field. Our findings suggest that during evidence integration, the population activity pattern does not simply scale at the target locations, as suggested by single-cell studies, but instead exhibits retinotopic shifts due to competitive dynamics during a deliberation period before transitioning to a commitment phase.

Our mechanistic circuit model demonstrates how competitive neural circuit dynamics induce the activity patterns representing the two choices to retinotopically shift toward one another during deliberation. This process reduces the influence of evidence on the decision process as the competition becomes more asymmetric (Fig. 4B). As one choice wins and its bump of activity grows under self-excitation, marking the commitment phase, the influence of evidence on the decision process is largely lost. This aligns with previous findings that later evidence contributes less to decision-making than earlier evidence [Huk and Shadlen, 2005, Wong et al., 2007, Stine et al., 2023]. It was proposed that this occurs as circuit self-excitation takes over from sensory evidence in driving neural activity [Wong et al., 2007], but without considering a dynamic shift in the neurons representing a choice and its gradual effect on the influence of evidence.

Previous work has identified signals from outside LIP that may influence termination of evidence accumulation. A time-dependent, evidence-independent urgency signal has been postulated to reduce the threshold to trial termination with increasing time [Hanks et al., 2014, Drugowitsch et al., 2012, Thura and Cisek, 2017]. A burst in superior colliculus has been shown to signal reaching of threshold, and suppressing superior colliculus prolongs the integration of evidence as revealed in behavior and LIP activity [Stine et al., 2023]. Such external signals might allow flexible adaptation of the processes described here, for example to control the speed-accuracy trade-off [Bogacz et al., 2010].

The model makes key predictions that we verify in the data. First, it predicts that the choice selectivity of individual neurons changes as the deliberation phase goes on, in a way that depends systematically on the retinotopic position of their response field. This finding may explain reports of heterogeneous response profiles of LIP neurons in similar tasks [Huk et al., 2017, Park et al., 2014], and may add insights into the origin of mixed selectivity observed in cognitive tasks more generally [Fusi et al., 2016]. Second, it predicts a behavioral effect: the slower the trial, the more the monkey undershoots the target, due to an increasing shift in neural activity toward the retinotopic center along with LIP’s established role as a topographic saliency map guiding saccadic eye movements [Bisley and Goldberg, 2010].

A recent study analyzing the same dataset found that confidence—defined as the probability that a decision is correct—can be linearly decoded from the activity, at the decision’s end, of the population of LIP neurons that represent the contralateral choice target [Zylberberg and Shadlen, 2025]. It is possible that the trial-to-trial variability across these neurons, which allows decoding, corresponds to the changing retinotopic location of the activity bump at the end of the decision. Greater retinotopic movement reflects greater competition between the two options, which should correlate with lower confidence.

We directly study trajectories in neural state space to determine the decision manifold, without prior assumptions about its structure. This revealed its curved, locally two-dimensional structure with one dimension representing reaction time, and the transition from deliberation to commitment. These results depend on our geometric characterization of neural trajectories, using arc length rather than time. This, along with the organization of the manifold by reaction time, was crucial to allow comparison of points in different trajectories at similar stages of the decision process (similar arc length and reaction time), regardless of the absolute time at which they occurred. An alternative method to achieve this is time warping (Supp. Fig. 2; Williams et al., 2020), but this does not allow single-trial analysis.

Other studies have assumed, and inferred, latent decision variables determining neural activities. [Genkin et al., 2023] found that a one-dimensional decision variable and decision manifold could largely explain neural activity in the primate dorsal premotor cortex (PMd) during a reaction-time task with two difficulty levels and two choices. The adequacy of their 1D assumption may stem from a narrower distribution of reaction times due to limited task difficulty levels, lower recording yields which could hinder detection of distributed representations, or fundamental differences in the functional roles of PMd vs. LIP. Luo et al. [2024] observed a transition from deliberation to commitment in a fixed-delay perceptual decision-making task in rodents. Such a transition in a fixed-delay task, representing the time the animal decides, could be the equivalent of the reaction time in a reaction-time task, whereas we have found such a transition well before the reaction time. They found the dynamics could be characterized by a 2D latent decision variable, specifying a flat 2D space in which neural activity lies, but the decision manifold within this space was close to one-dimensional. They modeled the transition using an extended drift-diffusion model (DDM) with an unspecified commitment mechanism, while we find it naturally emerges from intrinsic circuit dynamics. A previous circuit model pre-allocates distinct neural populations to encode each choice [Wong and Wang, 2006], effectively reducing the system to two latent decision variables – the activity level representing each choice – and thus a flat, 2D space. In contrast, the shift of activity in our model results in a curved non-planar decision manifold, smoothly modulating the contribution of individual neurons to the decision process depending on the strength of competition.

The DDM model of evidence integration successfully explains many aspects of noisy evidence integration, both psychophysical [Ratcliff, 1978, Bogacz et al., 2006, Ratcliff and McKoon, 2008] and neural [Gold and Shadlen, 2007, Shadlen and Kiani, 2013]. Our findings extend our understanding of neural evidence accumulation in two ways. First, previous work found that a linear sum of the activities of LIP T_in_ neurons directly represents the integrated evidence [Roitman and Shadlen, 2002, Huk and Shadlen, 2005, Churchland et al., 2011, Steinemann et al., 2024]. Thus, integration would represent movement in a single direction in neural state space. However, we found that integration of evidence occurs along a curved, two-dimensional manifold that results from shifting bumps of activity on the visuotopic map. The two points of view are not incompatible: the single direction might be the “readout”, observed by other areas, of the more complex neural evidence integration process. Indeed, in our model, which replicated psychophysical as well as neural findings, the decision process was terminated when a readout along a single dimension – a fixed linear sum over neurons surrounding a given target –exceeded a threshold. The second extension is that the population transitions to a commitment phase in which evidence is no longer integrated, and that this transition can occur hundreds of milliseconds before the decision’s report (*e*.*g*., see Supp. Fig. 6). In contrast, the DDM predicts that evidence is integrated until the moment a threshold is exceeded. Again, our results need not be incompatible with previous studies demonstrating DDM-like neural behavior [*e*.*g*., Steinemann et al., 2024, and references therein], because these studies, which sought to minimize effects of a decision bound on DDM signatures, were restricted to early times. Thus, the transition to commitment that we see might occur at times beyond these analyses. Indeed, there has long been an appreciation that evidence presented toward the end of a decision is often not incorporated [Huk and Shadlen, 2005, Wong et al., 2007, Stine et al., 2023, van den Berg et al., 2016, Resulaj et al., 2009]. Note that both of these extensions hold if we limit our analysis to T_in_ neurons (Supp. Fig. 4).

Previous studies have shown that significant variability in the activity patterns of a subset of LIP neurons that are highly selective for a target can be explained by a model in which single neurons do not reflect the integration of evidence but instead undergo a single, step-like change in activity within a trial [Latimer et al., 2015, Zoltowski et al., 2019]. Our analysis shows a sharp, synchronized, step-like transition from deliberation to commitment on slower trials (*e*.*g*. note the relatively sharp-in-time transitions in average single-trial behavior in Supp. Figs. 2G,6). This might account for at least some of the variability explained by a stepping model. Nevertheless, we also show that this step-like activity on the longest trials is preceded by a prolonged deliberation period in which, in the model, noisy evidence integration occurs, driving movement in the uncertainty direction. The stepping studies characterized neurons as steppers or non-steppers, but we have shown that neuronal trajectories can vary depending on reaction time and retinotopic response field locations (Fig. 5B,C), which may condition how well a neuron’s behavior is explained by a stepping model. Future analyses of stepping behavior at the population level should take these factors into account, and analyze whether population-synchronized steps occur and, if so, whether or how they relate to the deliberation/commitment transition revealed here.

Our analysis revisits the existing conceptual framework of the neural basis of decision-making, suggesting a central role for distributed and dynamic population activity. Our mechanistic circuit model emphasizes the role of neural competition between evidence alternatives in driving shifts in the decision-engaged neural populations, along with a transition from sensory-driven deliberation to circuit-dominated commitment. In sum, we present a unified mechanistic and population-dynamic picture of the emergence of a cognitive process from distributed network activity.

## Acknowledgements

We thank Larry Abbott, Stefano Fusi and Natalie Steinemann for helpful discussions. M.M. and K.M. were supported by the Gatsby Charitable Foundation (GAT3708), the Kavli Foundation, the NSF (DBI-1707398), and the NIH (U01 NS108683, R01 EY029999, U19 NS107613, RF1 DA056397). G.S. was supported by the NIH (T32 EY013933, F31 EY032791) and a Simon’s foundation postdoctoral fellowship. M.S. was supported by the Howard Hughes Medical Institute, the NIH (R01NS113113), the Grossman Center for the Statistics of the Mind at Columbia University, and the AFOSR (21RT0878).

## Author contributions

MM conceived, designed, and conducted the research presented in this study, with consistent input from and conversation with KM and input from GS and MS. MM wrote the original draft. MM, KM and GS revised and edited the manuscript, with input from MS. GS collected the data analyzed here in the lab of MS.

## 1 Differential Geometry in the Context of Decision-Making

Differential geometry offers a powerful framework for understanding the structure of the neural state space involved in cognitive processes like decision-making. In this section, we introduce fundamental concepts in differential geometry and illustrate how they relate to decision-making in the brain.

### 1.1 Manifolds and Intrinsic Dimensionality

A *manifold ℳ* is a mathematical structure that, while possibly complex globally, locally resembles Euclidean space ℝ*m*. More formally, an *m*-dimensional manifold is a topological space such that, for each point *p*∈ *ℳ*, there exists an open neighborhood *U* ⊂ ℳ containing *p* and a smooth map *φ* : *U →* ℝ^*m*^, where *φ* is a homeomorphism between *U* and an open subset of R^*m*^. This guarantees that, near any point, the manifold behaves like Euclidean space ℝ^*m*^.

The number *m* is called the *intrinsic dimensionality* of the manifold and indicates the number of independent coordinates required to describe the system locally. Even though the global structure may be complex, the manifold locally behaves like a subset of ℝ^*m*^.

Example 1: The 2D Sphere *S*^2^

Consider the surface of a 2-dimensional sphere *S*^2^ of radius *r*, embedded in 3-dimensional Euclidean space ℝ^3^. The equation of the sphere is given by:

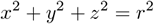

Here, *S*^2^ is embedded in ℝ^3^, but its *intrinsic dimension* is 2. Locally, we can describe the surface using two parameters, typically spherical coordinates (*θ, ϕ*), where *θ* ∈ [0, *π*] is the polar angle and *ϕ* ∈ [0, 2*π*] is the azimuthal angle. Thus, two coordinates are sufficient to describe any point on the sphere, even though the sphere is embedded in a higher-dimensional space.

Around each point on *S*^2^, the local structure can be mapped onto an open subset of ℝ^2^, making *S*^2^ a 2-dimensional manifold.

Example 2: The 2D Torus *T* ^2^

The 2-dimensional torus *T* ^2^ is another example, formed as the product of two circles:

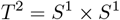

Each circle *S*^1^ is parameterized by an angular coordinate *θ* ∈ [0, 2*π*], so the torus can be described by two angular coordinates (*θ*_1_, *θ*_2_), where both *θ*_1_ and *θ*_2_ range from 0 to 2*π*. The intrinsic dimension of the torus is 2, since locally it resembles ℝ^2^, even though it has a more complex global structure.

While the torus can be embedded in ℝ^3^, its intrinsic dimensionality remains 2, reflecting its local 2-dimensional structure.

The intrinsic dimension of a manifold captures its fundamental degrees of freedom, and is independent of the dimension of the space in which it is embedded. For example, a curve *γ*, which is a 1-dimensional manifold, can be embedded in R^3^, but its intrinsic dimension remains 1. Similarly, the sphere *S*^2^ and torus *T* ^2^ are both intrinsically 2-dimensional, even though they can be embedded in spaces of higher dimension.

#### Isomap: Projection to Lower Dimensions

The Isomap algorithm is a widely-used technique for reducing the dimensionality of data while preserving the manifold’s intrinsic geometry. It seeks to discover the intrinsic structure of a high-dimensional dataset by constructing a lowdimensional embedding that best preserves the geodesic distances between points along the manifold.

Formally, let 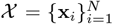 be a set of data points sampled from an underlying manifold ℳ embedded in a higher-dimensional Euclidean space ℝ^*n*^. The goal of Isomap is to find a lower-dimensional embedding 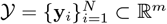 that approximates the geodesic distances on ℳ, as opposed to the Euclidean distances in ℝ^*n*^.

To achieve this, Isomap follows these steps: 1. Neighborhood Graph Construction: A neighborhood graph is built by connecting each point **x**_*i*_ to its nearest neighbors based on Euclidean distance. 2. Geodesic Distance Estimation: Geodesic distances between all pairs of points are estimated by finding the shortest paths on the neighborhood graph. 3. Multidimensional Scaling (MDS): The geodesic distance matrix is then used in classical MDS to find the low-dimensional embedding *Y*.

Unlike traditional linear methods such as Principal Component Analysis (PCA), which minimizes distortion in Euclidean space, Isomap preserves distances along the manifold itself. This is crucial for uncovering the true underlying structure of the data when the embedding space is significantly higher-dimensional than the manifold’s intrinsic dimension.

#### Estimating Intrinsic Dimensionality with Reconstruction Error

One of the advantages of Isomap is that it allows us to estimate the intrinsic dimensionality of the manifold by analyzing the reconstruction error. After applying Isomap to project the data into various dimensions *m* = 1, 2, 3, …, we can compute the reconstruction error, which quantifies how well the low-dimensional embedding preserves the manifold’s structure.

To compute the reconstruction error in our Isomap approach, we used the scikit-learn implementation, which calculates reconstruction error based on kernel matrices rather than directly comparing geodesic distances on the manifold and Euclidean distances in the embedded space.

The reconstruction error is defined as the Frobenius norm between two kernel matrices: the kernel matrix of the original high-dimensional data *D* and that of the embedding space distances *D*_fit_. Specifically, given a distance matrix *D* for the input data and an embedding distance matrix *D*_fit_, the reconstruction error *E* is calculated as follows:

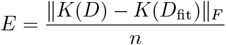

where *K*(*D*) is the centered kernel matrix for *D*, calculated as:

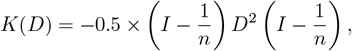

with *I* as the identity matrix, *n* the number of samples, and ∥· ∥ _*F*_ representing the Frobenius norm. The kernel matrix *K*(*D*) applies a centering operation to adjust for translation and scaling differences in the original and embedding spaces. By measuring this centered kernel difference, the error quantifies how well the low-dimensional embedding represents the relationships among data points present in the original space.

The reconstruction error in this approach does not capture the exact comparison of geodesic and Euclidean distances; rather, it uses a kernel-based method that indirectly reflects how well the embedding preserves local and global structures of the original manifold. Lower reconstruction error suggests that the embedding space closely approximates the true structure of the data.

By plotting the reconstruction error as a function of the embedding dimension *m*, we can observe a plateau when the embedding dimension matches the intrinsic dimensionality of the manifold. As the dimensionality increases, the reconstruction error decreases, but once the intrinsic dimension is reached, further increases in dimension lead to minimal improvement.

To robustly select the optimal number of neighbors *k*, which influences the geodesic distance calculation, we evaluated a range of values for *k* from 5 to 200 in 30 logarithmically spaced steps. For each choice of *k*, we ran the Isomap algorithm 30 times and computed the average reconstruction error. We then selected the value of *k* that minimized this average reconstruction error, which we report as our final reconstruction error. This approach provided a stable estimate, less sensitive to the inherent stochasticity of the algorithm.

Thus, Isomap provides a principled way to estimate the intrinsic dimensionality by identifying the point at which the reconstruction error stabilizes, while the averaging procedure ensures a more reliable choice of neighbors *k*. This method is particularly useful for real-world data where noise and measurement errors can obscure the true geometry of the underlying manifold.

### 1.2 Embeddings and Embedding Dimensionality

An *embedding* of a manifold is a smooth, injective map that allows the manifold to be realized as a subset of a higher-dimensional Euclidean space while preserving its geometric structure. Formally, given an *m*-dimensional manifold *M*, an embedding is a smooth map *f* : *M*→ ℝ^*n*^ such that *f* (*M*) is diffeomorphic to *M* and the differential map *df*_*p*_ has full rank for all points *p* ∈ *M*.

The *embedding dimensionality n* refers to the dimension of the ambient Euclidean space R^*n*^ in which the manifold can be embedded without distortion. Importantly, the intrinsic dimensionality of the manifold *M*, which is *m*, is independent of the dimensionality of the embedding space. The intrinsic dimension captures the local degrees of freedom of the manifold, whereas the embedding dimensionality accounts for the minimum dimension of the Euclidean space required to faithfully represent the manifold.

As an example, consider the 2-dimensional sphere *S*^2^, which is an intrinsically 2-dimensional manifold, meaning it can be locally parameterized by two coordinates (latitude and longitude). However, to embed the sphere in Euclidean space without distortion, we need at least a 3-dimensional space, ℝ ^3^. The intrinsic dimension of *S*^2^ is 2, but its embedding dimension in ℝ^3^ is 3.

Similarly, a torus *T* ^2^, which is a surface generated by revolving a circle around an axis in ℝ^3^, is an intrinsically 2-dimensional manifold, but can be embedded in ℝ^3^ or higher-dimensional Euclidean spaces.

#### Local Embedding Coordinates

When a manifold is embedded in a higher-dimensional space, it can be described locally in terms of *embedding coordinates*. These coordinates *ϕ* : ℝ^*m →*^ ℝ^*n*^ are the coordinates of the ambient space that locally describe the points on the manifold. For example, in the case of the sphere *S*^2^ embedded in ℝ3, we have three embedding coordinates (*x, y, z*) that describe points on the sphere, but only two of them are independent due to the constraint *x*^2^ + *y*^2^ + *z*^2^ = 1.

In general, a point on a manifold *M* embedded in ℝ^*n*^ can be described locally by *n* embedding coordinates. However, only *m* of these coordinates are independent, where *m* is the intrinsic dimension of the manifold. These local embedding coordinates provide a way to describe the manifold in the higher-dimensional space, while the intrinsic structure of the manifold remains independent of the embedding.

#### Whitney’s Embedding Theorem

A crucial result in differential geometry is Whitney’s Embedding Theorem, which provides a general answer to the question of how many dimensions are required to embed a manifold without distortion. The theorem states that any *m*dimensional smooth manifold can be embedded in ℝ2*m*. This means that for any manifold *M* of intrinsic dimension *m*, there always exists an isometric embedding in ℝ2*m*, but often the manifold can be embedded in even lower-dimensional spaces.

For example, Whitney’s theorem guarantees that a 2-dimensional manifold can always be embedded in R4. However, specific manifolds, such as the sphere *S*^2^, can be embedded in R3, which is a lower dimension than the general bound given by Whitney’s theorem.

### 1.3 Neural State Space and the Decision Manifold

The neural state space is a high-dimensional space where each axis corresponds to the activity of an individual neuron. If we consider a population of *n* neurons, the neural state space is then a Euclidean space R*n*, where each point in this space represents a particular state of neural activity. At any moment in time, the collective firing rates of the *n* neurons define a point in R*n*, and as these firing rates evolve over time, they trace out a trajectory in this space.

While the neural state space is typically high-dimensional (with *n*≫ 1), not all possible trajectories are meaningful for the behavioral task at hand. In the case of decision-making, the neural activity associated with this process is constrained to a much lower-dimensional structure within the high-dimensional neural state space. This lower-dimensional structure is what we refer to as the *decision manifold*. In the context of decision-making, we found that this manifold has an *intrinsic dimensionality* of two, meaning that locally, the space of decisions is spanned by just two degrees of freedom, even though the embedding (neural) space is of much higher dimension.

We identify the two essential variables that capture the key dynamics of the decision process. The first variable, which we refer to as **strength of the evidence**, reflects the relative confidence or bias toward one choice over another. This corresponds directly to the external information provided to the decision-maker, such as the clarity or coherence of sensory stimuli. Behaviorally, we can infer this strength of evidence from measures like reaction time, with stronger evidence typically leading to faster responses, and weaker or ambiguous evidence leading to slower decisions.

The second variable represents the **evolution of the decision over time**. This describes how the decision unfolds as information is processed and accumulated, beginning from the stimulus onset and continuing through the decision phase until the final response, such as a saccadic eye movement. This progression can be thought of as a temporal trajectory that represents the dynamic process of integrating evidence. When parameterized by arc length, this variable allows us to capture the path the decision follows in terms of gradual shifts in cognitive state toward one of the decision boundaries. Together, these two variables—strength of evidence and decision evolution—are not merely abstract constructs but are tightly linked to the observable behavior of the subject. They provide a behavioral and cognitive framework for understanding the structure of the decision-making process. By representing the decision manifold in terms of these two dimensions, we gain insights into the underlying neural mechanisms and their role in driving observable choices. This is particularly important because these variables anchor the manifold in meaningful, interpretable terms, revealing how the brain transitions between different states as it forms and executes a decision.

Although the decision manifold has an intrinsic dimension of two, it is embedded in a much higher-dimensional neural state space. This embedding reflects the fact that even though decisions are made based on two main variables, these variables are realized through the coordinated activity of a large number of neurons. The *embedding dimensionality* of the decision manifold is the minimum number of dimensions in the neural state space required to faithfully represent the geometry of the decision manifold without distorting its structure.

In real-world data, especially in neuroscience, estimating the embedding dimensionality of a manifold is challenging due to noise, measurement errors, and the variability inherent in biological systems. This complexity can obscure the true geometric structure of the data, making it difficult to distinguish between noise and meaningful variation. A commonly used method to approximate the dimensionality of the embedding space is the *participation ratio*.

The participation ratio provides an estimate of the number of significant dimensions in the data by analyzing the eigenvalue spectrum of the covariance matrix. Given the covariance matrix **C** of neural activity, with eigenvalues *λ*_*i*_ (which correspond to the variance explained by each principal component), the participation ratio is computed as:

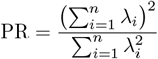

Here, *n* represents the number of eigenvalues (dimensions). The participation ratio provides an estimate of how many dimensions carry significant variance in the data, discounting small contributions from noise. A low participation ratio indicates that the data is effectively confined to a lower-dimensional subspace, even though it exists in a higher-dimensional embedding space.

For neural data, the participation ratio can give insight into how many dimensions are necessary to capture the main variance in neuronal activity, as opposed to noise. This becomes crucial when trying to understand the structure of the neural state space, where the decision manifold is embedded.

#### Submanifolds of Decisions

A further observation is that the decision manifold is composed of two disconnected *submanifolds*, corresponding to the two alternative choices. Each of these submanifolds reflects the trajectory of the decision process for one particular choice, and they are disconnected because each trial leads to a binary outcome. Therefore, the decision manifold is formed by these two disjoint components:

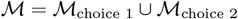

where each ℳ_choice_ is a submanifold representing one of the two possible decisions. These submanifolds are disconnected in the sense that no continuous path exists between the two choices on the manifold.

### 1.4 Parametric Curves in High-Dimensional Space

A **parametric curve** or **trajectory** in a high-dimensional space is a smooth mapping from a real parameter (typically time) to the points in a space of higher dimension. Formally, let ℳ represent a manifold embedded in an *n*-dimensional Euclidean space, ℳ*n*. A parametric curve on this manifold is a function:

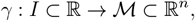

where *I* is an interval of the real line representing the parameter (which could be time, arc length, etc.), and *γ*(*t*) =

**x**(*t*) = (*x*_1_(*t*), *x*_2_(*t*), …, *x*_*n*_(*t*)) describes the coordinates of the curve in the embedding space at each time *t* ∈ *I*.

This curve *γ*(*t*) is said to be **smooth** if all of its component functions *x*_*i*_(*t*) are continuously differentiable. The image of *γ*(*t*) represents the path traced by the curve as the parameter *t* varies over the interval *I*.

Example:

Consider the simple case of a 2-dimensional plane, ℝ2, where we define a parametric curve by:

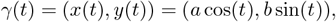

where *a* and *b* are constants. This defines an ellipse, a well-known example of a parametric curve in two dimensions. For each value of *t*, the function *γ*(*t*) provides a specific point (*x*(*t*), *y*(*t*)) on the ellipse.

Interpretation in the Context of Decision-Making:

In the context of the decision-making manifold, each parametric curve can represent the trajectory of a single trial in the high-dimensional space of neural activity. As the decision-making process unfolds, the brain moves through different configurations of neuronal firing rates, which can be captured as a trajectory in this high-dimensional space. The parameter *t* could represent the passage of time during the decision process, and the curve *γ*(*t*) would describe the path the brain takes through this space from stimulus onset to the decision response.

By studying the geometry of such trajectories, we can gain insight into how the decision evolves over time. In particular, we can analyze how the brain transitions between different states, how the evidence accumulates, and how the final decision is executed, all of which are captured by the curve’s shape and properties in the embedding space.

#### Natural Parametrization (Arc-Length Reparametrization)

In general the parameter *t* does not correspond to any intrinsic property of the curve but is simply an arbitrary parameter such as time. A more natural way to parametrize a curve is through its **arc length**, which measures the actual distance traveled along the curve. This gives rise to what is known as **arc-length parametrization** or **natural parametrization**.

Let *γ*(*t*) be a parametric curve in ℝ^*n*^. The arc length *s* of the curve from *t*_0_ to some *t* is defined as:

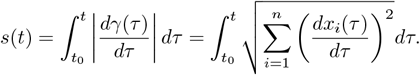

This formula computes the cumulative distance traveled along the curve from *t*_0_ to *t*, where 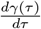 is the velocity vector of the curve and 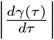 is its magnitude, i.e., the speed along the curve at parameter value *τ*.

To reparametrize the curve in terms of arc length, we introduce a new parameter *s*, such that *s*(*t*) is the arc length from some initial point. The parametric curve in terms of *s* is then written as:

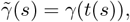

where *t*(*s*) is the inverse function of *s*(*t*). When a curve is parametrized by its arc length, the speed along the curve is always constant and equal to 1, i.e.,

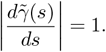

Example:

Consider the ellipse from the previous example, parametrized by *t*:

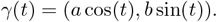

The arc length of this ellipse from *t*_0_ = 0 to any *t* can be computed as:

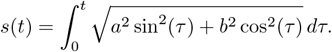

The curve can then be reparametrized by arc length *s*, which gives a more natural description of the distance traveled along the ellipse.

#### Interpretation in the Context of Decision-Making

In the context of decision-making trajectories, arc-length reparametrization allows us to express the decision process in terms of how much progress has been made along the decision trajectory, rather than in terms of time. We further normalize each trial’s arc-length to run from 0 to 1 to better compare equivalent points across nearby trajectories. This is particularly useful when a set of trials have a large heterogeneity in their reaction times, as the arc length represents the intrinsic evolution of the decision process. The evolution of a decision is thus measured in terms of how far the system has progressed along the manifold, providing a time-invariant way of analyzing the decision dynamics.

By analyzing decision trajectories in terms of their arc length, we can compare decisions that take different amounts of time but follow the same geometric path in neural space. This allows us to better understand the geometry of decision-making as it unfolds on the neural state space.

#### Distance Between Trajectories

Given two parametric curves *γ*_1_(*s*) and *γ*_2_(*s*), both parametrized by normalized arc-length *s* ∈ [0, 1], we can define a *distance* between the two trajectories as the integral over their normalized arc-length of the absolute difference between the two curves. This distance, denoted by *d*(*γ*_1_, *γ*_2_), is given by:

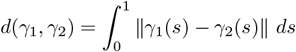

where ∥·∥ is the Euclidean norm in the ambient (embedding) space, and *s* is the normalized arc-length, ranging from 0 to 1. The normalization ensures that regardless of the absolute length (in Hz) of the curves in neural space, we are comparing them over a common scale of progress along their paths. This is particularly useful in decision-making contexts, where trajectories may differ in duration and total neural length but can still be meaningfully compared based on their normalized progression.

Intuitively, this distance measures the total cumulative deviation between the two paths, capturing both how far apart they are at each point and how their shapes differ over time. If the trajectories are identical, the distance will be zero, while larger distances indicate more significant divergence between the two curves at different points along their paths.

#### Tangent Vector to the Curve

For a parametric curve *γ*(*t*) in ℝ^*n*^, the **tangent vector** to the curve at any point is the vector that represents the direction of the curve at that point. Mathematically, the tangent vector **T**(*t*) is defined as the derivative of the curve with respect to its parameter:

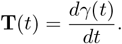

This vector points in the direction of the curve at parameter value *t*, and its magnitude represents the speed at which the curve is being traversed. When the curve is reparametrized in terms of arc length *s*, the tangent vector becomes a unit vector, meaning its magnitude is always 1:

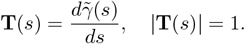

#### Interpretation in the Context of Decision-Making

In the context of decision trajectories on the decision manifold, the tangent vector represents the direction in neural state space of the decision process at any given moment. As a decision evolves from evidence accumulation to action selection, the tangent vector captures the direction of this progression. This indicates the instantaneous contribution of single neurons in LIP to changes in the evidence integration process.

#### First-Order Curvature

The *curvature κ*(*t*) of a smooth parametric curve *γ*(*t*) in ℝ^*n*^ is a measure of how quickly the curve is deviating from being a straight line. Mathematically, it is defined as the magnitude of the rate of change of the unit tangent vector **T**(*t*) with respect to the arc length *s*:

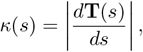

where **T**(*s*) is the unit tangent vector along the curve:

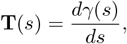

and *s* is the arc-length parameter.

In practice, especially when dealing with discretized data or noisy curves, the above method can lead to numerical instability. A more robust approach to estimate the local curvature at a point in the curve uses the ratio of the arc length to the chord length for a small segment of the curve centered at that point. The curvature is computed as:

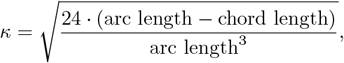

where the arc length is the length of the segment, and the chord length is the straight-line distance between the segment’s start and end points in high dimensional euclidean space. This method avoids the direct use of derivatives, which are sensitive to noise, and instead uses geometric properties of the curve.

Curvature gives a local description of how much the curve bends at any point. A straight line has zero curvature everywhere, meaning its tangent direction does not change. A circle has constant non-zero curvature, indicating that its tangent vector continuously changes direction at a fixed rate. The larger the curvature, the sharper the curve bends at that point.

#### Interpretation in Decision-Making

In the context of decision-making on the neural state-space manifold, curvature can describe how rapidly the decision process changes direction over time. A trajectory with low curvature indicates a smooth, steady decision process, while regions of high curvature suggest rapid changes in the decision trajectory, possibly due to conflicting or fluctuating evidence during the decision process. Thus, curvature can quantify how neural dynamics, evidence accumulation, and external stimuli shape the trajectory on the decision manifold. This measures how much the state changes smoothly from one set of neurons to another.

#### Tortuosity

Tortuosity is a measure that captures the complexity or “wiggleness” of a curve. Unlike curvature, which describes the local bending of the curve, tortuosity provides a characterization of how a curve sharply and momentarily deviates from a constant curvature. For example, a circle has (possibly high) constant curvature but zero tortuosity because it does not “wiggle” in any complex way. However, irregular or highly convoluted curves may exhibit high tortuosity.

Mathematically, tortuosity is often defined using higher-order derivatives of the curve’s position. However, for a more intuitive and practical definition that can be robustly computed for experimental data, we define tortuosity as the squared ratio of the derivative of curvature with respect to arc length to the curvature itself:

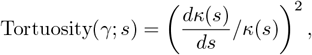

where *κ*(*s*) is the curvature as a function of arc length *s*.

Tortuosity provides a way to quantify the complexity of a curve beyond just its local bending. A curve with high tortuosity could be one that loops or twists frequently, even if its individual segments have low curvature. This is in contrast to curvature, which captures only the rate of turning at a specific point.

For example: A straight line has both zero curvature and zero tortuosity. A circle has constant curvature but zero tortuosity, as its curvature does not change. A highly irregular or zigzagging curve could have low curvature at most points but high tortuosity due to frequent changes in curvature.

#### Tortuosity in Neural State Space

In the context of neural state-space trajectories, tortuosity can be used to describe how erratic or convoluted the decisionmaking process is. If the decision manifold trajectory has high tortuosity, it may indicate a more complex decision-making process, with frequent shifts in neural dynamics due to changing evidence or internal fluctuations. For example, a high-tortuosity trajectory could correspond to a scenario where the decision process is influenced by multiple competing sources of evidence, causing the decision trajectory to take a more convoluted path through the state space, or it may indicate the effect of internal or external sources of noise irrelevant to the decision process.

##### Higher-Order Curvatures in High-Dimensional Space

To fully describe the geometry of a curve beyond its local bending, we define higher-order curvatures using Gram determinants. Higher-order curvatures provide insight into how the curve twists and evolves in space beyond the firstorder curvature. These curvatures are calculated from a sequence of derivatives of the curve in high-dimensional space and capture increasingly complex geometric properties.

#### Gram Determinants and Curvatures

Let *γ*(*s*) ∈ ℝ^*n*^ be a smooth curve parameterized by arc length *s*, where *n* is the dimensionality of the embedding space. We define the first, second, and higher-order derivatives of the curve with respect to *s*:

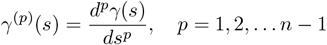

To compute the curvatures, we first form the Gram determinant *G*_*p*_(*s*), which is the determinant of the matrix composed of the first *p* + 1 derivatives of *γ*(*s*). Specifically, for each point *s*, the Gram determinant is:

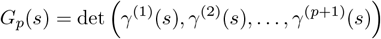

The higher-order (or generalized) curvatures *κ*_*p*_(*s*) are then computed using these Gram determinants. The first curvature *κ*_1_(*s*) (the standard curvature) is:

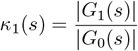

For higher-order curvatures, such as the second curvature *κ*_2_(*s*) (torsion) and beyond, the formula is:

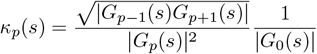

The first curvature measures how much the curve fails to be a straight line at a given point. Similarly the second curvature (torsion) measures the failure of the curve to be locally constrained to a plane at that point.

#### Fundamental Theorem of Space Curves in High Dimensions

The Fundamental Theorem of Space Curves provides a crucial result in differential geometry that fully characterizes a curve in Euclidean space up to rigid transformations. Specifically, it states that a space curve is completely determined by its curvature and torsion (or higher-order curvatures in higher-dimensional spaces) if given an initial position and an initial frame of reference.

#### Statement of the Fundamental Theorem

For a smooth curve **r**(*s*) ∈ ℝ^*n*^ parameterized by arc length *s*, the curve is uniquely determined, up to Euclidean motions (rotations and translations), by its sequence of curvatures *κ*_1_(*s*), *κ*_2_(*s*), …, *κ*_*n−*1_(*s*) and an initial reference frame (**r**(0), **r**^*′*^(0), …, **r**^(*n−*1)^(0)).

In other words, if two curves share the same curvatures *κ*_1_(*s*), *κ*_2_(*s*), …, *κ*_*n−*1_(*s*), and have the same initial frame, then they are identical up to rigid motions in space.

#### Interpretation in Neural State Space

In the context of neural state space, the Fundamental Theorem suggests that the dynamics of a neural trajectory—such as the evolution of a decision-making process—can be fully described by its curvatures. This implies that, given the curvatures of the neural trajectories, one can reconstruct the entire decision-making process in the high-dimensional neural space. This principle allows to compare different trajectories based only on their shape disregarding a decision’s orientation or location in state space, i.e. independently of what set of neurons are encoding the trajectory.

### 1.5 Tangent Spaces of Embedded Surfaces

A fundamental concept in differential geometry is the notion of the tangent space at a point on a manifold. In this subsection, we will define the tangent space to an embedded surface, explain how to compute it, and show its relevance to understanding neural computations on the decision manifold.

#### Tangent Space at a Point on an Embedded Surface

Given a smooth manifold ℳ embedded in a higher-dimensional Euclidean space ℝ^*n*^, the tangent space at a point *p* ∈ *ℳ* is the set of all possible directions in which one can move along the surface from *p*. Formally, it is the vector space of the derivatives of all smooth curves on ℳ that pass through *p*.

Let **x**(*t*) be a smooth parametric curve on the surface ℳ with **x**(0) = *p*. The tangent vector to this curve at *p* is given by the derivative:

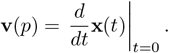

The collection of all tangent vectors to curves passing through *p* forms the tangent space at *p*, denoted by *T*_*p*_ ℳ. It is a linear subspace of ℝ^*n*^ whose dimension is equal to the intrinsic dimension of the manifold ℳ.

For example, for a 2-dimensional surface embedded in ℝ3, the tangent space at a point *p* is a 2-dimensional plane passing through *p* that is tangent to the surface.

#### Basis of the Tangent Space: Local Coordinate Representation

The tangent space *T*_*p*_ℳ can be spanned by a set of basis vectors that describe the directions of possible movement in the neighborhood of *p*. Ifℳ is parametrized by local coordinates (*x*_1_, *x*_2_, …, *x*_*m*_), where *m* is the intrinsic dimension of the manifold, the basis vectors of the tangent space are given by the partial derivatives of the embedding functions **F**(*x*_1_, *x*_2_, …, *x*_*m*_):

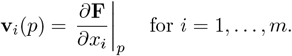

These vectors span the tangent space at *p* and form the basis for any tangent vector at that point.

#### Tangent Space and Neural Computation

In the context of neural state space, the neural decision manifold is a 2-dimensional surface embedded in the highdimensional space of collective neuronal activity. The local tangent space at each point on this manifold is defined by two key vectors:

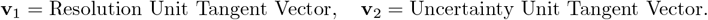

These vectors represent the interpretable directions in which the neural system can change in response to the decisionmaking process. Specifically:

- **v**_1_ (the resolution vector) represents the direction in the neural state space that is tangent to the average decision trajectory, specifically capturing changes along the arc-length coordinate of the decision path.
- **v**_2_ (the uncertainty vector) represents the direction in the neural state space orthogonal to the resolution vector. It lies within the plane defined by the resolution vector and the tangent to the local reaction-time coordinate and points toward faster reaction times.

The tangent space at any point on the decision manifold represents the computationally relevant directions for the neural system. These directions either capture the gradual progression of the decision process from stimulus onset to saccade (resolution direction) or account for sudden deviations that influence the resulting reaction time, perhaps signaling momentary evidence (uncertainty direction). Together, these vectors describe how the neural population dynamics evolve to encode decisions through evidence accumulation.

#### Orthogonal Directions and Noise

In addition to the computationally relevant tangent space, the neural trajectories may also extend in off-manifold directions, which are orthogonal to the decision manifold in the high-dimensional neural state space. These directions, denoted as off-manifold projections, capture irrelevant noise that does not contribute directly to the decision-making process.

Noise in neural activity is not uniformly distributed. Some noise may project onto the tangent space and influence the decision-making process, while other noise projects into the orthogonal off-manifold space and has no direct effect on decision computation. The key distinction between these two types of noise is critical for understanding how decision accuracy and reaction time are affected.

By quantifying the projections of noise onto the tangent space and the off-manifold space, we can determine how much of the observed variability in neural activity is behaviorally relevant. The off-manifold noise is considered irrelevant to the decision process, and its influence can be mitigated.

The decision manifold’s tangent space thus provides a powerful framework to separate signal from noise in neural data. This allows us to precisely quantify the effects of noise on decision computations and understand the robustness of neural decision-making in the presence of stochastic fluctuations and trial-to-trial variability.

Why Tangent Spaces Matter for Neural Computation

Understanding the structure of the tangent space at each point on the decision manifold is crucial for several reasons:

- It helps identify the dimensions along which neural computations of evidence integration and decision evolution occur.
- It allows for a decomposition of the neural signal into behaviorally relevant and irrelevant components.
- It provides a means to quantify the effect of noise on decision-making, allowing us to distinguish between computational noise (in the tangent space) and irrelevant fluctuations (off-manifold).

In summary, the tangent space provides a local linear approximation of the manifold at each point, where the neural system performs its decision-making computations. The off-manifold direction captures irrelevant noise that does not contribute directly to decision computation. By studying the structure of tangent spaces and their relationship to noise, we can better understand the computational principles underlying decision-making in the brain.

### 2 Single Trial inference Using AutoLFADS

To accurately characterize the geometry of the decision-making state space, it is essential to obtain smooth representations of neural spiking activity on a single-trial basis. Raw spiking data often contains high levels of noise and irregularities, making it difficult to infer the underlying firing rates of neurons across trials. To address this, we employed the AutoLFADS (Latent Factor Analysis via Dynamical Systems) deep learning technique [1]. AutoLFADS allows us to extract smooth, trial-by-trial estimates of the neural firing rates from population-level recordings.

AutoLFADS is a generative recurrent neural network model that extends the LFADS framework by introducing variational autoencoders and dynamical system priors. The model takes raw spiking data from a population of neurons and infers latent dynamical factors that evolve over time, which govern the neural activity. These factors are used to generate smoothed firing rate estimates that reflect the neural activity driving the behavior, while effectively reducing noise. The network also employs a controller to model external inputs that might influence neural dynamics, thus improving the model’s robustness to noise and unmeasured variables.

Neural recordings, particularly at the single-trial level, exhibit significant variability due to external noise and biological factors. Traditional trial-averaging methods are often insufficient for recovering the true underlying dynamics, especially in complex decision-making tasks where each trial may follow different trajectories in neural state space. AutoLFADS provides a probabilistic framework to decompose noisy, high-dimensional neural data into smooth, interpretable latent states, which are essential for reconstructing accurate neural dynamics in cognitive processes. This approach enables us to capture trial-to-trial variability while preserving the temporal structure of the data.

#### Data Preparation

We processes and run AutoLFADS for each session independently. We divided the trials into training and validation datasets based on their choices (left or right) and coherence levels, ensuring a well-balanced contribution of trials from all conditions such that the training set comprised two-thirds of the trials, with the remaining one-third used for validation. Spike data from each trial were binned into time windows spanning from the onset of the dot stimulus to the completion of the saccade. The spike trains for each neuron were aligned to the start of the trial and binned into 10 ms intervals.

In addition to the neural data, external inputs were created to reflect the duration and stimulus direction of the trial. Two trial-independent binary vectors, one for each stimulus direction, were constructed to indicate when the dot motion stimuli were present on the screen (1 for dots on, 0 for dots off). This provided context to AutoLFADS about when stimulus-related neural activity occurred, assisting in the reconstruction of firing rates for each trial. An implementation without these external input generates qualitatively similar results, but the external input helps with sharp boundaries (sudden start and stop of stimulus presentation).

#### Training Parameters

Once the data were prepared, they were passed to AutoLFADS for training, we use the online NeuroCAAS framework for training [2]. The training and validation data consisted of both the binned spike data and the corresponding external inputs. AutoLFADS was configured with a set of hyperparameters optimized for our dataset. The primary LFADS parameters found after optimization by AutoLFADS include the following:

**Table 1.** Optimal AutoLFADS model parameters used for training of the example session.

| Parameter | Value | Description |
| --- | --- | --- |
| batch_size | 128 | Number of trials per training batch. |
| encod_data_dim | 138 | Number of neurons in the input data. |
| encod_seq_len | 225 | Sequence length of the encoded data. |
| ext_input_dim | 2 | Dimension of external inputs (left and right dots). |
| fac_dim | 40 | Dimensionality of factors . |
| gen_dim | 128 | Dimension of the generator. |
| ic_dim | 64 | Dimension of the initial conditions. |
| kl_co_scale | 0.79 | KL divergence scaling for controller output. |
| kl_ic_scale | 0.0068 | KL divergence scaling for initial conditions. |
| l2_con_scale | 274.87 | L2 regularization scaling for controller. |
| lr_init | 0.001 | Initial learning rate. |
| lr_decay | 0.95 | Learning rate decay factor. |

AutoLFADS was trained using these parameters to infer smooth firing rate estimates from the noisy binned spike data, with the external inputs providing task-related context to guide the reconstruction of the neural population activity.

### 3 Sequence Classifier for Reaction Time Quantile Prediction

To predict reaction time quantiles from the sequence of neural activity data, we implemented a classifier that combines Long Short-Term Memory (LSTM) networks and Fully Convolutional Networks (FCN). This architecture captures patterns in the sequential data, which were reparametrized by arc length. The classifier was built in PyTorch and trained using PyTorch Lightning for streamlined model optimization and evaluation.

#### Model Architecture

The model, *Multivariate LSTM-FCN* [3], is designed for multivariate sequence classification. It processes multiple input features simultaneously, capturing patterns across these features to improve classification performance. The model is based on an augmented version of the univariate LSTM-FCN network, extended to handle multivariate data effectively. By leveraging both LSTM and fully convolutional networks (FCNs), the model captures complex interactions within the sequences of multiple neural features, offering a flexible and powerful architecture for decision quantile prediction.

- **LSTM**: The LSTM network processes the multivariate sequence data, capturing dependencies across multiple features. A three-layer LSTM with 512 units per layer was used to model these dependencies. The output from the last LSTM layer represents the processed feature interactions over the full sequence.
- **FCN**: In parallel, the sequence is passed through a fully convolutional block. This block consists of three 1D convolutional layers, with kernel sizes of 8, 5, and 3 and channel dimensions of 128, 256, and 128. Each layer captures interactions across the multivariate input features at different scales. The network includes a squeeze-and-excitation block to enhance feature representation, boosting classification accuracy.
- **Multivariate Integration**: The outputs from the LSTM and FCN are concatenated, combining both sequential feature dependencies and local feature patterns. This allows the model to efficiently handle multivariate data by considering both sequential evolution and inter-feature relationships.
- **Fully Connected Layers**: The combined outputs are passed through two fully connected layers, with 100 units in the hidden layer and softmax activation in the final layer to output the predicted reaction time quantile.

#### Training and Optimization

The classifier was trained using the processed neural data, split into training (60%), validation (20%), and test (20%) sets. Stratified splitting ensured balanced representation of reaction time quantiles across sets.

We used the Adam optimizer with a learning rate of 0.001. To manage the learning rate decay, we employed a learning rate scheduler (StepLR) that reduced the learning rate by a factor of 0.9 every 10 epochs. Early stopping was implemented based on validation loss, with a patience of 5 epochs to prevent overfitting.

**Table 2.** Hyperparameters used for the Multivariate LSTM-FCN model.

| Parameter | Value |
| --- | --- |
| LSTM Hidden Dimension | 512 |
| LSTM Layers | 3 |
| FCN Channels | 128, 256, 128 |
| Kernel Sizes (FCN) | 8, 5, 3 |
| Dropout Rate | 0.5 |
| Batch Size | 64 |
| Learning Rate | 0.001 |
| Optimizer | Adam |
| Scheduler | StepLR (factor: 0.9) |
| Early Stopping Patience | 5 epochs |
| Max Epochs | 100 |

The model was trained for 50 iterations, and those achieving a test accuracy of at least 35% were aggregated for further analysis. Confusion matrices were generated to evaluate classification performance across reaction time quantiles. Confusion matrices highlight the ability of the model to differentiate between adjacent quantiles effectively.

### 4 Smooth Representations of Scalar Variables vs. Reaction Time and Arc Length

To analyze how trial-averaged trajectories evolve and how single-trial deviations influence cognitive processing, we developed a method to compute a smooth representation of scalar variables (e.g., neuron firing rates) as a function of both arc length and reaction time. This is achieved by applying cross-validated non-parametric regression (LOWESS: locally weighted scatterplot smoothing) followed by interpolation for each arc length point, allowing us to visualize the relationship between reaction time and variables across trials.

For each analysis, trials corresponding to a specific choice are filtered from the dataset. We include correct and incorrect trials in all analyses with no distinction. For a given arc length value, scalar values (e.g., neuronal activity) across all trials are plotted against the reaction time (RT). Optionally, large outliers based on reaction time are removed using a predefined threshold of 50 ms to filter trials with significant RT gaps.

#### LOWESS Smoothing and the CDF Approach

To ensure that the smoothing process is optimal, we perform cross-validation on the LOWESS smoothing fraction parameter, which controls the span of the smoothing window. The smoothing process can be performed either directly on the reaction times or after transforming the reaction times using their cumulative distribution function (CDF). The CDF transformation standardizes the reaction times by mapping them into a uniform distribution, potentially improving the performance of the LOWESS smoothing in cases where reaction times are non-uniformly distributed.

When the CDF method is applied reaction times are first transformed into a uniform distribution using their cumulative distribution function (CDF). LOWESS smoothing is then applied on the transformed reaction times, followed by interpolation to map the smoothed values back to the original reaction time space. This method is useful when the original reaction time distribution is extremely skewed, and uniformizing it via CDF can enhance the accuracy of the smoothing step, effectively implementing a locally adaptive smoothing value that depends on the trial count around a specific reaction time.

#### Cross-Validation Process

For each arc length point, the following procedure is applied:

- **Training and Validation Splitting**: The reaction times (RTs) and scalar values from all trials are split into training and validation sets using an 80/20 split. The training set is used for smoothing, and the validation set is used to evaluate the model.
- **LOWESS Smoothing**: For each value of the smoothing fraction (*frac*) in the range [0.1, 0.9], we apply the LOWESS smoothing algorithm to the training data, generating smoothed values across the RTs. This step can be applied either directly on the reaction times or after the CDF transformation.
- **Interpolation**: Cubic interpolation is applied to the smoothed training data, mapping the smoothed values to the validation RTs.
- **Mean Squared Error Calculation**: We compute the Mean Squared Error (MSE) between the predicted and true validation values for each fraction value. The fraction that minimizes the MSE across the validation set is selected as the optimal smoothing fraction.

This cross-validation process ensures that the LOWESS fraction is tuned to avoid overfitting or underfitting the data, resulting in an optimal representation of scalar variables across reaction times. The optimal fraction is used to smooth the scalar values for all trials at each arc length point, producing a trial-averaged representation.

Once the optimal smoothing fraction is determined, we apply the LOWESS algorithm to the entire dataset, using the optimal fraction to compute smoothed values of the scalar variables as a function of reaction time. After smoothing, we perform cubic interpolation to generate a consistent RT axis across all trials. The final output is an array of smoothed and interpolated values of the scalar variable across all arc length and reaction time points.

**Table 3.** Parameters used for cross-validated LOWESS smoothing.

| Parameter | Value |
| --- | --- |
| Fraction Range for LOWESS | 0.1 - 0.9 |
| Number of RT Points | 101 |
| Test Size (Cross-Validation) | 0.2 |
| Interpolation Method | Cubic |
| RT Gap Threshold | 50 ms |
| CDF Option | Optional (uniformizing reaction times) |
| Data Filter | Optional (outlier removal) |

### 5 Visuotopic mapping of choice selectivity

The monkeys performed a visually instructed delayed saccade task [4], where a target was presented at a pseudo-random location in the visual field. The fixation point remained visible for a variable delay, ranging from 0.4 to 1.5 seconds, depending on the monkey’s training. After the delay, the fixation point was extinguished, signaling the monkey to make a saccade toward the previously shown target.

#### Neural Response Field Calculation

To compute the firing rate maps (or response fields) of neurons during the target presentation, we aligned the spike data to the target onset. For each trial, we counted the number of spikes (*N*_spikes_) in a window centered at time *t* after the target was presented. The window had a duration of 2*w*, where *w* is half the window size. The firing rate (*FR*) for each neuron in trial *i* was computed as:

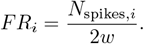

The firing rates were computed across different target positions (*x, y*) in the visual field and then grouped to compute the average firing rate per neuron across all trials for each target position.

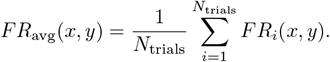

To address potential occupancy biases due to the non-uniform distribution of target locations in the visual field, we applied Gaussian smoothing to the computed response fields divided by the occupancy (the sum of the gaussians centered at the target locations). This process ensures that each neuron’s firing rate is represented on a consistent spatial grid, reducing the bias that would otherwise arise from uneven coverage of the visual space.

We applied Gaussian smoothing with a kernel of standard deviation *σ* = 3 degrees of visual angle. The kernel was applied over a grid spanning from `mindeg` = —15^°^ to `maxdeg` = 15^°^, using a resolution of *N*_side_ = 101 grid points per axis. This approach regularized the firing rate estimates by distributing activity over adjacent grid points based on their proximity in the visual field.

The smoothed response field at each grid point is calculated as a weighted average of the firing rates within the Gaussian window. This can be expressed as:

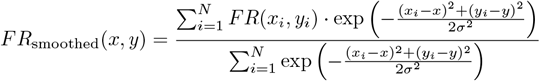

where:

- *FR*(*x, y*) is the resulting firing rate at visual grid position (*x, y*),
- *FR*(*x*_*i*_, *y*_*i*_) is the firing rate at target position (*x*_*i*_, *y*_*i*_),
- *σ* is the standard deviation of the Gaussian kernel,
- The numerator represents the Gaussian-weighted sum of firing rates, and
- The denominator normalizes this sum, ensuring the weights sum to 1.

This process provided a consistent spatial representation of neural activity, effectively reducing the bias introduced by irregular target placements while preserving the spatial structure of the neural response fields.

To determine the optimal time shift (*t*) and window size (*w*) for computing the neural response fields, we computed Moran’s I as a measure of spatial clustering. Moran’s I quantifies the degree of spatial autocorrelation, indicating how similar nearby grid points in the response field are in terms of neural activity. It is defined as:

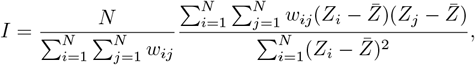

where:

- *N* is the number of grid points,
- *Z*_*i*_ is the firing rate at location *i*,
- 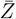 is the mean firing rate across all locations,
- *w*_*ij*_ is the weight, modeled as a Gaussian:

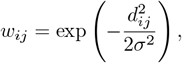

where *d*_*ij*_ is the Euclidean distance between grid points *i* and *j*, and *σ* is the same value used in the spatial smoothing.

Moran’s I was computed for different values of *t* and *w*, ranging from 0.1 to 0.3 seconds for *t* and 0.005 to 0.2 seconds for *w*, with a step size of 0.01. The goal was to maximize spatial clustering, indicating that neurons showed spatially consistent firing rate patterns across the visual field. Moran’s I provides a way to enhance the spatial information in neural response fields by identifying regions of the visual field where neurons consistently respond similarly.

The parameter grid search provided a clear maximum. The optimal window parameters were found to be *t* = 0.150 seconds and *w* = 0.085 seconds, which maximized the Moran’s I value, indicating the highest degree of spatial autocorrelation in the response fields.

The table below summarizes the key parameters used in computing the neural response fields:

**Table 4.** Summary of parameters used in response field computations.

| Parameter | Value |
| --- | --- |
| Time shift ($t$) | 150 ms |
| Window size ($w$) | 85 ms |
| Smoothing $\sigma$ | 3 dva |
| Minimum firing rate | 10 Hz |
| Spatial grid size | 101x101 pixels |
| Range of grid (mindeg) | -15 degrees |
| Range of grid (maxdeg) | 15 degrees |

#### Mapping choice selectivity to the visual field

To map any scalar variable associated with the neural population, such as choice selectivity, onto the visual field, we utilized each neuron’s response field as a spatial filter. This allowed us to interpolate neural properties based on the spatial organization of neurons’ receptive fields. For instance, the choice selectivity of neurons for specific conditions (e.g., slow or fast trials) was projected onto a grid representing the visual field.

The scalar variable *P*_*i*_ corresponding to neuron *i* was mapped to the visual field by weighting the neuron’s response field *Z*_*i*_ (a smoothed representation of the firing rate map) according to *P*_*i*_. Specifically, for each grid point, the property map *M* (*x, y*) was computed as a weighted sum of the properties across all neurons:

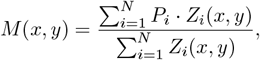

where *Z*_*i*_(*x, y*) represents the Gaussian-smoothed response field of neuron *i* at location (*x, y*).

To enhance the contrast between the peaks of activity and regions of noisy, spontaneous firing outside the neurons’ response fields, we exponentiated the normalized response fields before applying the weighted sum. This step increases the contrast between regions with high neural selectivity and regions with low or irrelevant background activity, improving the representation of spatial information. The exponentiation operation was performed as follows:

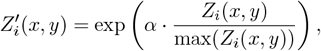

where *α* is a scaling factor (set to 18 in our case) that controls the level of contrast enhancement.

This procedure allowed us to generate more informative maps of neural properties, such as choice selectivity, by minimizing the influence of irrelevant background noise and focusing on regions where neurons showed meaningful spatial tuning.

The final smoothed maps of neural properties, such as choice selectivity, exhibited clear spatial organization, providing insight into how neurons’ selectivity varies across the visual field. The optimal window parameters for computing response fields were *t* = 0.150 seconds and *w* = 0.085 seconds, which were chosen based on the maximum spatial clustering as measured by Moran’s I.

### 6 Mechanistic Model of Decision-Making on the LIP Retinotopic Sheet

To investigate the neural mechanisms underlying decision-making in the lateral intraparietal area (LIP), we constructed a spatiotemporal model that simulates the interaction between excitatory and inhibitory neurons distributed across a two-dimensional retinotopic sheet. Notably, in contrast to previous models, the model does not explicitly encode choice in neuron subpopulations but instead represents neural population activity dynamically as it integrates noisy evidence information in favor (or against) the choice targets in response to distributed sensory input.

Neurons in the model represent positions in the visual field, with connectivity strength decaying as a function of distance in retinotopic space. Excitatory neurons exhibit a rectified linear response and inhibitory neurons have a rectified quadratic response.

Neurons receive external mean inputs modeled as spatial activity bumps at the locations of the contra and ipsi targets. These inputs are linearly modulated by the signed coherence of the stimulus, representing sensory evidence for each target. Furthermore, excitatory neurons receive spatially correlated pink noise, capturing variability in sensory input and neural activity.

#### Continuous Model Equations

The dynamics of excitatory (*r*_*e*_(**x**, *t*)) and inhibitory (*r*_*i*_(**x**, *t*)) neural populations at each location **x** = (*x, y*) on the retinotopic sheet are described by the following differential equations:

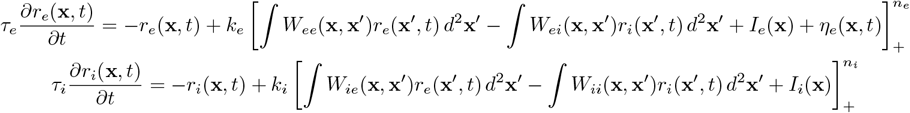

where:

- *r*_*e*_(**x**, *t*) and *r*_*i*_(**x**, *t*) are the activities of excitatory and inhibitory neurons at retinotopic position **x** and time *t*,
- *W*_*αβ*_(**x, x**^*′*^) represents the synaptic weight from neuron type *β* at position **x**^*′*^ to neuron type *α* at position **x**,
- *I*_*e*_(**x**) and *I*_*i*_(**x**) are the external inputs for excitatory and inhibitory neurons,
- *η*_*e*_(**x**, *t*) denotes spatially correlated colored noise, and
- [·]_+_ denotes the rectified linear unit (ReLU) nonlinearity applied to the inputs.

The synaptic weights follow a Gaussian decay with retinotopic distance:

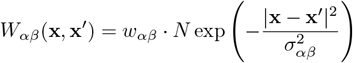

where *w*_*αβ*_ represents the synaptic strength between neurons of types *α* and *β*, and *σ*_*αβ*_ controls the spatial spread of the connections.

The integrals in the equations above sum the inputs from neurons across the retinotopic sheet. The normalization constant *N* ensures that the total connectivity strength to a a postsynaptic neuron equals *w*_*αβ*_. This is done numerically to correct for boundary effects in the simulation sheet.

#### External Inputs

The mean external inputs *I*_*e*_(**x**) and *I*_*i*_(**x**) are taken to be equal in these simulations. They are modeled as two activity bumps centered at the positions of the contralateral (contra) and ipsilateral (ipsi) targets. These activity bumps are described by Gaussian functions:

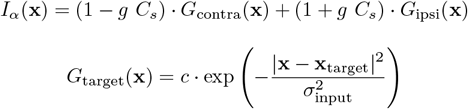

where *σ*_input_ controls the spatial extent of the target inputs, *c* is a scaling factor (the Gaussians are not normalized), *g* is a gain factor, and the input amplitude is modulated by logarithmically spaced values of the signed coherence *C*_*s*_.

Each location **x** on the retinotopic sheet experiences spatially correlated noise *η*_*e*_(**x**, *t*), modeled as an OrnsteinUhlenbeck (OU) process in time. The noise is correlated across space by convolving spatially independent OU processes at each spatial point with a Gaussian filter.

The temporal evolution of spatially independent OU noise traces 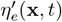 is governed by :

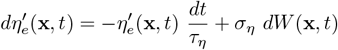

where *τ*_*η*_ is the noise time constant, *σ*_*η*_ is the noise amplitude, and *W* is the temporal Wiener process. Numerically we integrate the following Euler-forward update rule

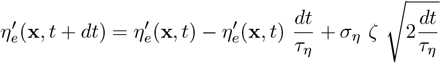

where at each timestep and independently for each location we sample *ζ* from a normal distribution with zero mean and unit variance. The spatially correlated noise *η*_*e*_(**x**, *t*) is constructed through convolution with a Gaussian kernel:

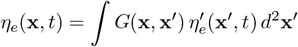

where 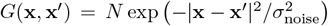 is the spatial correlation kernel. The normalization factor *N* implements ∫ Ω *G*^*2*^ (**x, x**′) *d*^*2*^ **x**′ = 1, where Ω is the boundary of the simulation sheet. The normalization ensures resulting processes have the same variance and time correlation while correcting for boundary effects numerically.

Parameters used in the model are given in Table 5. In experiments, the two targets are in opposite visual fields and thus primarily represented in opposite hemispheres. The inhibitory to excitatory connections are artificially long (*σ*_*ei*_ is twice the inter-target distance) to represent suppression between target activities in the two hemispheres, known to occur in frontal eye fields [5] and which we postulate similarly occurs in LIP, but whose anatomic basis we did not try to accurately model. This choice is not critical to the model; for example, setting *σ*_*ei*_ = 10 and multiplying the *w*’s by 0.65 gives similar results.

**Table 5.**
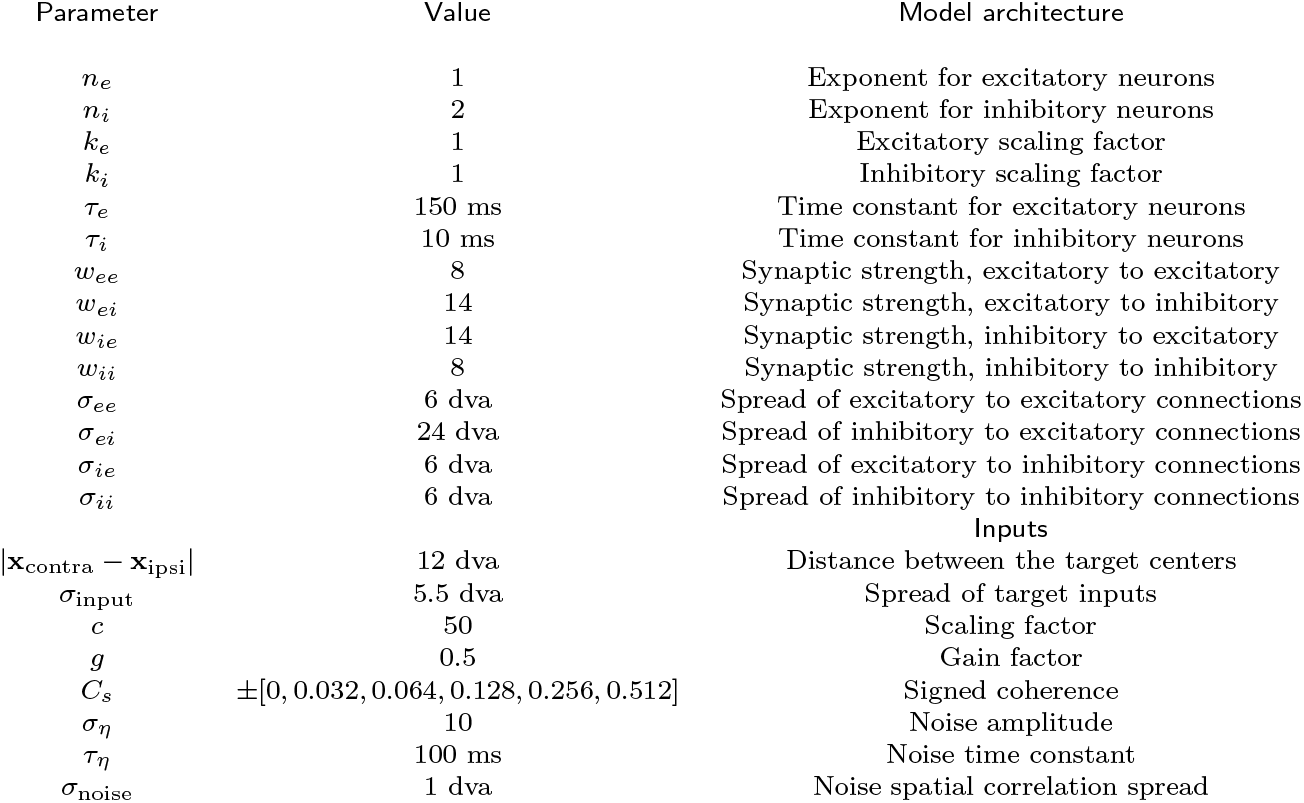
Summary of parameters used in the LIP retinotopic model.

#### Simulation and Post-Processing

We simulate the model on a 31× 31 spatial grid, representing a retinotopic sheet with a side length of 30 degrees of visual angle (dva). Each simulation runs for 2 seconds with a time resolution of *dt* = 0.5 *ms*, consistently reaching a stable steady state. The initial conditions influence the timing of early dynamics, shifting the distribution fo reaction times, we use *r*_*e*_(*t* = 0) = *r*_*i*_(*t* = 0) = 10.

For post-processing, the first 300 ms of each simulation (corresponding to the period where neural activity has not yet separated for different choices) is discarded. A decision is declared when the difference in the average neural activity around the two target locations reaches a threshold of 70 Hz. The activity around each target is averaged within a radius of 3 dva, and the time at which this threshold is crossed is recorded as the reaction time for that trial.

The neural activity is smoothed using a Gaussian kernel with a temporal standard deviation of *σ* = 50 ms and subsampled to a time resolution of 10 ms. Additionally, trials with reaction times in the top and bottom 3% are excluded from further analysis to eliminate outliers.

### 7 Local resolution- and uncertainty-driving input directions

Our aim is to analyze the influence of momentary evidence on neural activity along the Resolution and Uncertainty projections on the contra submanifold of neural state space. The contra submanifold is defined by trials with a contra-choice. We constructed local area patches across this submanifold, dividing trials into arc-length × reaction-time segments to capture the effect of the momentary evidence at distinct points in the decision manifold. This is important because of the manifold non-linear and curved representation, suggesting that at different stages during the decision process evidence might have a different influence on the cognitive directions within the manifold.

1. Reaction Time Mapping: To control for varying trial counts across RT, we used a CDF-based reparameterization. Each trial’s RT value was mapped into a 0-1 RT range using the cumulative distribution function of RTs. This allowed us to construct RT windows that are evenly populated across trials.
2. Arc × RT Patches: We defined each patch as a region in the arc × RT space, where arc specifies a segment of the arc length of the neural trajectory, and RT defines a specific range of RTs. This method enabled us to isolate patches that contain localized segments of neural trajectories for trials falling within a specific RT range. Each patch captures a subset of arc length for trials within a given RT range, allowing us to focus on a small, spatially coherent region of the curved submanifold.
3. Contra Submanifold: Only trials within the contra-choice manifold were included. Each patch hence represents a localized segment of the contra submanifold, allowing us to study how evidence shapes neural fluctuations across similar conditions.

For each arc × RT patch, we analyzed how neural activity fluctuates in response to evidence-related fluctuations in the input. The input for each patch, *X*_*i*_(*t*), representing the external input to each excitatory neuron on trial *i* at arc length *t*, is defined as a matrix of shape (trials × *A* × *N* ^2^), where the arc-length segment of the patch is discretized into *A* evenly-spaced points, and *N* is the retinotopic (pixel) grid size so that *N* ^2^ is the number of excitatory neurons. To study how input fluctuations influence activity fluctuations in the Resolution and Uncertainty directions, we applied two steps of normalization to the input:

1. Mean Centering: The neural input *X*(*t*) was first mean-centered within each patch:

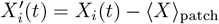

where ⟨*X*⟩ _patch_ is the mean input across trials and arc-length points in the patch. This provides a mean input vector of shape *N* ^2^ for each patch, representing the average evidence state in the patch capturing the noise-free mean input determined by the coherence level.
2. L2 Normalization: We normalized each trial’s input by its L2 norm over the pixel dimension to standardize fluctuation magnitude:

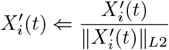 This normalization means that we are assessing the effect of the direction of deviations from the mean input on the Resolution and Uncertainty projections, irrespective of the size of the deviations.

#### Fitting the local resolution- and uncertainty-driving input directions

To model the influence of fluctuations in the input 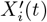 on the Resolution and Uncertainty projections within each arc × RT patch, we fit local input directions to capture the sensitivity of each projection to momentary evidence. These filters capture the directional influence of the momentary evidence on neural fluctuations.

1. Local Input Directions: For each patch, we fit separate local input directions for the Resolution (**w**_Res_) and Uncertainty (**w**_Unc_) projections, where each direction **w**_Res_ and **w**_Unc_ is of shape (N × N). These directions were trained to capture how the moment-to-moment fluctuations in the input evidence for each patch align with fluctuations of neural activity in the Resolution and Uncertainty directions.
2. Prediction Model: For all trials and arc-points within a patch, we used the following models to predict the resolution and uncertainty components of the trial’s unit tangent vector at that arc length:

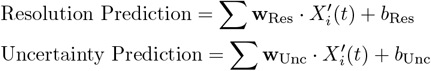

where *b*_Res_ and *b*_Unc_ are the biases for Resolution and Uncertainty, respectively. Each patch has independent biases, representing the baseline activity level in each direction.
3. Loss Functions and Optimization: To fit **w**_Res_ and **w**_Unc_, we minimized a loss function:
  a. Mean Squared Error (MSE) Loss: This component minimizes the squared error between the predicted and actual projections for all trials and arc-length points within a patch:

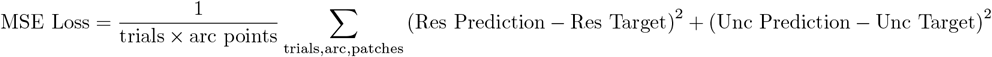
  b. Spatial Derivative Regularization: A second-order spatial derivative penalty was applied to the evidence directions to reduce high-frequency noise in retinotopic (pixel) space. This helps avoid overfitting to fine-grained fluctuations that may not be signal-relevant:

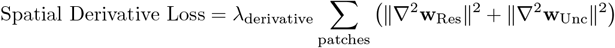

where ∇^2^ denotes the second-order spatial derivative (Laplacian), smoothing the evidence directions while preserving meaningful trends.

The total loss function is thus:

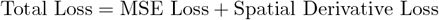

The local input direction parameters were optimized across multiple training epochs to ensure that each patch’s local direction captures the spatial effects of momentary evidence along the Resolution and Uncertainty projections.

#### Evidence Modulation Measure

We defined a global evidence direction in retinotopic (pixel) space, denoted **evdir**. This direction is computed as a contrast vector between contra and ipsi target locations:

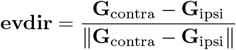

where **G**_contra_ and **G**_ipsi_ are the Gaussian functions that determined the deterministic input centered at the contra and ipsi target locations, respectively, with equal amplitudes, and the denominator normalizes this vector to unit length. This unit target-based direction measures the spatial influence of the evidence for the contra alternative across the retinotopic grid, serving as a consistent baseline to which we compare the local resolution- and uncertainty-driving input directions in each arc × RT patch.

To quantify the impact of momentary evidence on neural fluctuations, we computed an evidence modulation measure for each patch, capturing the alignment of the local input directions **w**_Res_ and **w**_Unc_ with the global evidence direction **evdir**:

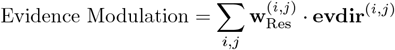

where *i, j* index the pixel grid. This measure reflects the strength with which the global evidence for the contra alternative actually influences the decision-making process at different stages and levels of difficulty.

**Table 6.** Summary of parameters used in local patch construction and filter fitting.

| Parameter | Value | Description |
| --- | --- | --- |
| Arc Window Size | 0.1 | Fraction of arc length range per window |
| Arc Window Step | 0.1 | Fractional step for arc length window movement |
| Arc Points | 10 | # of discrete arc-length points for each arc length window |
| RT Window Size | 0.1 | Fraction of RT CDF range per window |
| RT Window Step | 0.1 | Fractional step for RT window movement |
| $\lambda_{\text{derivative}}$ | 0.001 | Relative weight for the derivative loss |

This comprehensive approach reveals how Resolution and Uncertainty projections encode evidence and respond to momentary fluctuations, providing insights into neural mechanisms underlying decision-related activity.

## Supplementary Material 1: LFADS Performance in Fitting Spike Count Data

To assess the performance of LFADS (Latent Factor Analysis via Dynamical Systems) in capturing the underlying neural activity, we compared the smoothed neural activity inferred by LFADS with the observed spike counts from single trials. LFADS provides us with estimates of the underlying firing rates at single-trial and single-neuron resolution, which cannot be directly compared to the raw spike count data due to the inherent noise in spike trains. However, we can meaningfully compare the representations provided by LFADS by considering the information at the population level. Specifically, LFADS takes advantage of correlations across neurons to infer a low-dimensional latent representation that underlies the population activity. By leveraging this, we can compare how well LFADS fits single-trial population activity, and subsequently assess the inferred single-trial activity for each neuron using the population as a whole.

The benefit of this approach lies in the fact that neural population activity often exists on a low-dimensional manifold. This implies that the collective population rate contains distributed, repeated information across neurons, helping to average out the noise from individual neurons or trials. Thus, LFADS works to reduce noise in single-trial estimates by capturing the shared structure across the population. First, we show how well LFADS reconstructs single-trial activity when considering information from the whole population, using dimensionality reduction to compare inferred rates and actual data in this reduced space. This approach reveals the fidelity of LFADS in recovering smooth neural dynamics at the population level, which then informs the interpretation of single-neuron activity on a trial-by-trial basis.

### Comparison Measures

To quantify how well LFADS fits the spike count data at the single-trial level, we computed two key metrics. First, the squared relative error (SRE) was used to capture the divergence between the LFADS-inferred and observed data. For each trial and for each principal component (PC) of the spike data, we define the squared relative error as:

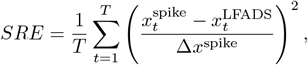

where 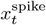 and 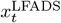 represent the time series of the principal component (PC) for spike count and LFADS data, respectively, and Δ*x*^spike^ = —max(*x*^spike^) − min(*x*^spike^) is the range of the spike data PC over time.

Second, we computed the Pearson correlation coefficient for each trial to assess the linear relationship between the inferred and Gaussian-smoothed (*σ* = 50 ms) observed spike data in PC space. For each trial and each PC, the Pearson correlation is given by:

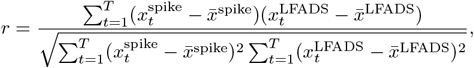

where 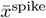 and 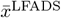 represent the mean of the spike data and LFADS data PCs, respectively.

### Performance Results

We computed both metrics for the first ten principal components of the spike count and LFADS data (Fig. 1B,D). The squared relative errors were computed as a percentage, with the median results for the first ten PCs reported as 3.7, 3.9, 3.9, 4.3, 4.4, 4.4, 4.6, 4.6, 4.6, 4.7 percent. The Pearson correlations for the first ten PCs, reflecting the linear fit between the spike count and LFADS data, yielded the following median values: 0.96, 0.93, 0.94, 0.83, 0.89, 0.63, 0.74, 0.44, 0.72, 0.52.

These results demonstrate that LFADS accurately captures the latent dynamics underlying neural activity at both the population and single-trial levels. The high Pearson correlations for the first several PCs, along with the relatively low squared relative errors, indicate a good fit between LFADS-inferred rates and the observed spike count data, particularly in the components capturing the most variance in the population activity.

### Average Decision Manifold Using Neuron Principal Components of Spike Data

Given the reasonable fit between LFADS and spike count data across the population, we attempted to recover the decision manifold directly from the spike data, using the principal components derived from the raw spike counts. The aim was to investigate whether similar geometric properties of the decision manifold—such as separability, alignment, and curvature—could be observed when analyzing the spike data in this reduced PC space (Fig. 1E-H). However, the inherent noise in the spike data, even after smoothing with a Gaussian kernel (*σ* = 50*ms*), presented significant challenges for performing single-trial analyses.

Evidence for the noisiness of the spike data can be seen in the higher relative errors and lower Pearson correlations that we computed for the first several principal components. Additionally, the participation ratio, a proxy measure for dimensionality, was found to be 23 for the smoothed spike data (Fig. 1C), indicating that many more components are required to capture the variance in the population. Specifically, 50 principal components were needed to explain 80% of the variance in the smoothed spike data, whereas only 3 principal components were sufficient to explain the same proportion of variance in the LFADS-inferred data. This contrast underscores the ability of LFADS to denoise neural activity and recover the underlying low-dimensional structure more effectively than the raw spike counts.

The spike data was smoothed with a Gaussian kernel with a standard deviation of 50 ms to reduce noise, but this was insufficient for capturing the organized fluctuations in relation to the decision manifold. The level of tortuosity, reflecting the curvature of trajectories in neural state space, was much higher in the spike data and was not clearly organized in local coordinates, suggesting a loss of alignment with the cognitive phases of deliberation and commitment. Furthermore, the off-manifold component—representing the fluctuations that cannot be explained by the manifold—reached up to 95% of the total fluctuation strength for slow trials, further highlighting the overwhelming influence of noise in the spike data. These observations lead us to conclude that the noise in the raw spike data, even after smoothing, obscures the true structure of fluctuations relative to the average manifold. As a result, while population-level analyses using principal components can provide some insight into the global geometric properties of the decision manifold, the noise in single trials makes it difficult to resolve finer details, such as those related to cognitive processing phases like deliberation and commitment.

### Choice Selectivity of Neurons Using Spike Data Averaged Over Trials

To assess how well LFADS fits the raw neural activity at the single-neuron level when averaged over trials, we divided the trials into reaction time quantiles for each choice. Specifically, we divided the data into 10 quantiles, with the top decile corresponding to the slowest trials and the bottom decile to the fastest trials. For each quantile, we reparametrized the trials by arc length and computed averages across trials for each neuron, separately for each choice. This provided four averaged datasets: slow-ipsilateral, slow-contralateral, fast-ipsilateral, and fast-contralateral, representing the neuronal dynamics for each condition.

We then compared the LFADS-generated neural activity and the smoothed spike count data by calculating the Pearson correlation coefficient for each neuron between these two datasets. The results of these comparisons showed that the LFADS data fits well with the raw spike count data. The median Pearson correlations were 0.92 and 0.89 for the slow contralateral and ipsilateral conditions, and 0.98 and 0.94 for the fast contralateral and ipsilateral conditions, respectively (Fig. 1I). However, the averaged slow decile did not fit as well, likely due to the increased noise in slow trials, which obscures the structure in the neural state space. This is particularly evident in the cognitive phases of deliberation and commitment, which are less clearly separated in the slow trials.

In the slow condition, the noise in single trials, particularly off-manifold noise, results in poor alignment of trials in arc length. This causes the average activity at earlier arc lengths to remain noisy, hiding the clustering of choice selectivity across neurons. To address this, we computed the choice selectivity for the last 10% of the arc length in the slow condition, as the neural activity at this point represents the commitment phase, which is more strongly aligned between trials. In contrast, the fast condition, where trials show a stronger commitment phase, allowed us to compute choice selectivity across the entire arc length.

The choice selectivity, calculated as (1 — Pearson correlation between contralateral and ipsilateral conditions)*/*2, was computed for each neuron in both the slow and fast conditions. We then plotted the distribution of differences in choice selectivity across neurons and observed that the changes in selectivity followed a clustering pattern consistent with the results obtained using LFADS in the main text. This demonstrates the robustness of our analysis in capturing cognitive changes at the single-neuron level (Fig. 1J-K).

**Supplementary Figure 1:**
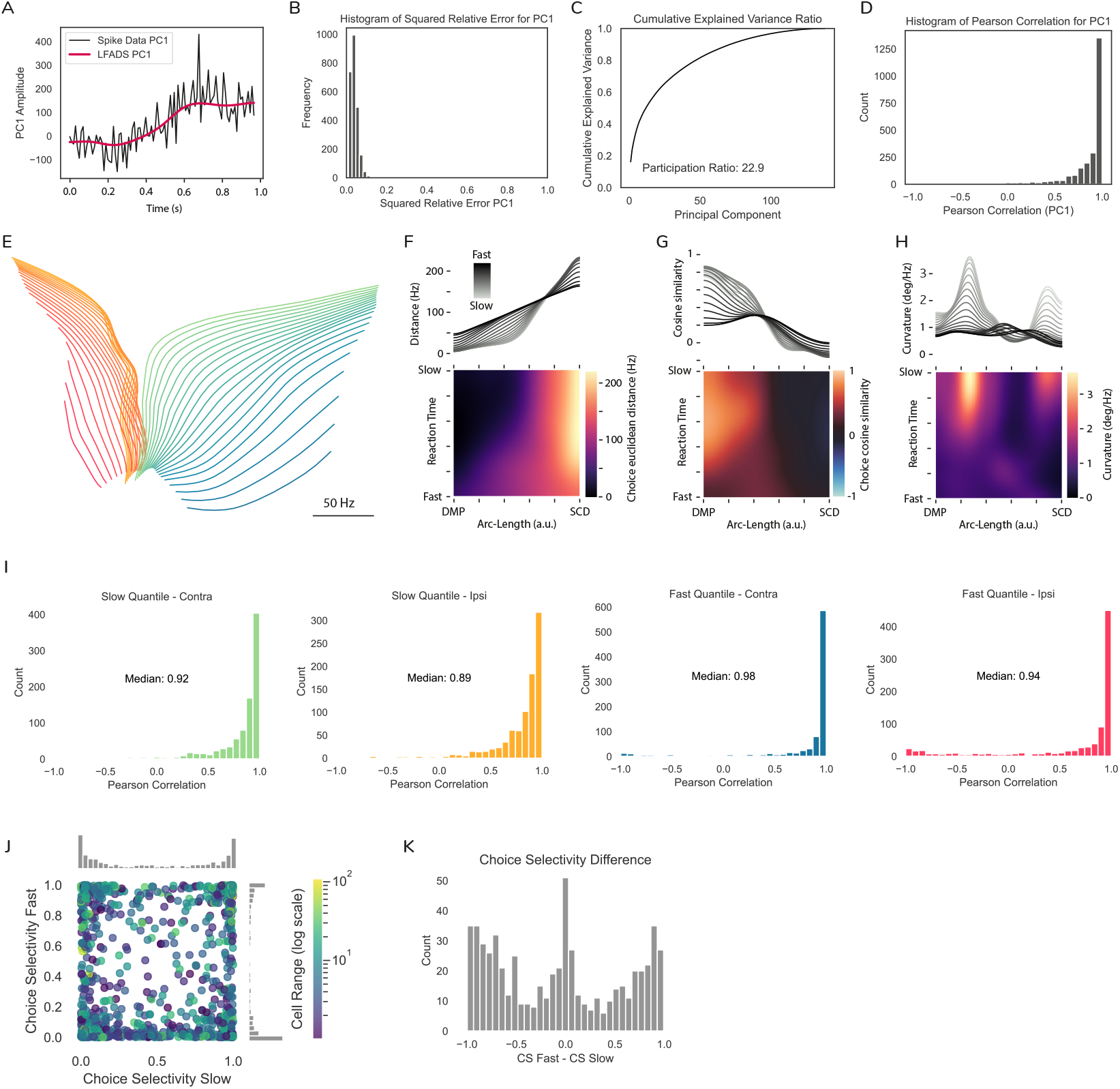
LFADS Performance in Fitting Spike Count Data. **Fig. 1 Comparison of LFADS-Inferred Rates and Raw Spike Count Data.** Panels (A) and (B) show results from PCA on the neuron dimension of the LFADS data, with the raw spike count data projected into the same PCA space. (A) Example single trial displaying the first principal component (PC1). The black trace represents the spike count data, and the red trace corresponds to the LFADS-inferred smooth firing rate for the same trial. This trial is the one closest to the median relative square error from panel (B). (B) Distribution of the relative square error between LFADS and spike count data for PC1 across trials. The text provides the median values for the first 10 PCs. Panels (C)-(H) use a Gaussian-smoothed version of the raw spike count data (kernel width *σ* = 50 ms). (C) Variance explained by a PCA performed on the smoothed spike count data. (D) Distribution of Pearson correlations between PC1 from LFADS and smoothed spike count data across trials. (E) Isomap projection of the average decision manifold using the first 50 PCs of the smoothed spike count data. (F)-(H) Geometric properties of the average manifold constructed from smoothed spike count data: separability (F), alignment (G), and curvature (H). (I-K) Both smoothed spike count and LFADS data were divided into 10 reaction time quantiles per choice. Trials from the top (slow) and bottom (fast) deciles were averaged for each choice, after reparametrizing trials by arc-length. (I) Distribution of Pearson correlations between LFADS and smoothed spike count data across neurons with a dynamical range of at least 1 Hz (N=894), shown for each average decile and choice. (J) Scatter plot of the choice selectivity for slow versus fast averages, illustrating the clustering of selectivity. (K) Distribution of the differences in choice selectivity between slow and fast conditions across neurons.

## Supplementary Material 2: Time warping

In this section, we applied a time-warping technique [1] to align neural activity across trials with varying reaction times, producing consistent trial-averaged representations of neural firing, as shown in Figure 2. Time-warping reduces variability in the timing of key task events, such as dot motion onset and saccade execution, while preserving the essential temporal and neural dynamics of the task. This approach allows us to study how neural activity evolves across different reaction time quantiles and choices, providing insights into the temporal and spatial dynamics of decision-making.

Figure 2A shows a spike raster plot for the slowest quantile of leftward choices. Before time-warping (left panel), spiking activity exhibits significant variability in the timing of key events, such as the onset of dot motion and saccade execution. After applying time-warping (right panel), this variability is minimized, yielding a more consistent alignment across trials. In Figure 2B, we see the trial-averaged activity of a representative neuron with a response field aligned with the left target. The time-warped activity (black trace) closely matches the peristimulus time histogram (brown trace), indicating that the time-warping method effectively smooths the neural activity into a coherent template while preserving key behavioral events like dot motion onset and saccade execution, as marked by diamonds and squares, respectively.

Figure 2C illustrates the progression of reaction times across quantiles, showing a near-linear increase in median reaction time from fast (Q1) to slow (Q10) quantiles. Blue and red lines represent leftward and rightward choices, respectively. Time-warping maintains the relative timing of key events, as seen in Figure 2D. While the median timing of these events remains consistent (left panel), the variability across trials is greatly reduced (right panel), resulting in a narrower distribution and greater synchronization of neural activity.

In order to connect this analysis to existing literature, we plot the average activity over time of neurons whose response fields are predominantly aligned with either the target in the contralateral visual field (TinC neurons) or the ipsilateral visual field (TinI neurons), see Figure 2G. Following the onset of dot motion stimulation, we typically observe a characteristic dip in neural activity lasting about 100ms, succeeded by an approximate 100ms ramp-up in activity. During this initial 200ms period, the responses do not exhibit a clear separation between choices. However, post-200ms from dot motion onset, the choices begin to distinctly diverge, and this separation appears to be reaction time-dependent, with slower quantiles showing a later separation in time between choices. Additionally, we give a first description of the corresponding neural activity within state space during these task stages (Figure 2H,I). The transition from the activity dip to the ramp is marked by a sharp directional change in state space, with the ramp itself extending around 150 Hz. This phase may represent some form of preparatory alignment prior to the integration of evidence. Around 200ms after the onset of dot motion, there is a pronounced directional shift and organization into an approximately two-dimensional surface in state space, with slower quantiles exhibiting a delayed separation between choices.

**Supplementary Figure 2:**
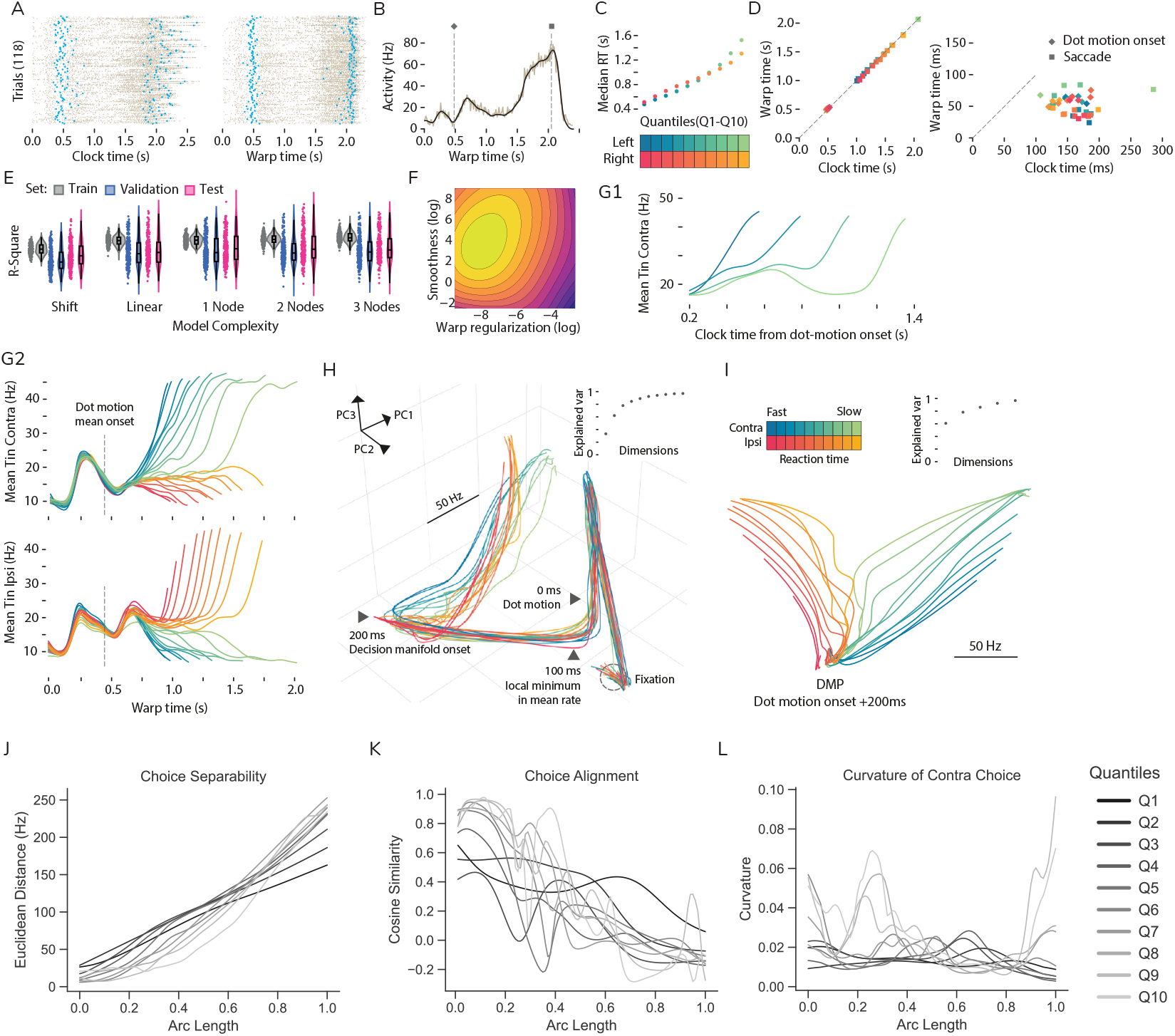
Time warping. **Fig. 2 Time Warping Technique for Consistent Trial-Averaged Neural Activity.** (A) Spike-time raster representation for trials within the slowest quantile (Q10) for leftward choices. The spiking activity, as observed in the experiment’s chronological order (left panel), demonstrates noticeable fluctuations during key task events: the initiation of dot motion stimuli and the saccadic action (illustrated by blue dots in the raster plot). Post time-warping, this variability in event timings is significantly minimized (right panel). (B) Average activity across trials for a representative neuron with a response field coinciding with the left target’s location. The time-warping method yields a smooth template representing the neuron’s typical activity pattern (depicted by the black trace). This closely aligns with the 10ms interval-based peristimulus histogram (represented by the brown trace). The diamond and square markers highlight the median timings of dot motion initiation and saccadic movement, respectively. (C) The median reaction time within each quantile exhibits a near-linear escalation corresponding to the quantile’s ranking. The blue and red color schemes represent choices towards the left and right targets, respectively. (D) Time-warping maintains the median timings of pivotal behavioral events (left panel), implying that neural activities in the warped timeframe mirror actual chronological neural activities. Conversely, the spread in behavioral event timings is considerably narrowed (right panel) as indicated by the diminished interquartile range of the distributions, pointing to a consistent synchronization across trials. (E,F) Model Selection and Validation: The model’s complexity is defined by its piecewise linear warping function, mapping chronological and warped time. The validation process spans over 200 trial and neuron configurations and 250 hyperparameter sets to select model complexity (E) and parameters like firing rate template smoothness and warping intensity (F), determined from the consolidated *R*^2^ values of the top quintile of models in the test set. (G-I) Visualization of time-warped quantile-averaged activity in both time and state space. (G) Displays the average activity of neurons with response fields aligned with either the contralateral (top panel; G1 shows 4 example trials) or ipsilateral (bottom panel) targets, differentiated by choices directed contra-laterally (blue) and ipsi-laterally (red). (H) Shows a PCA of the template population activity encompassing the entire experimental setup. (I) Highlights a PCA focused solely on activity beginning 200ms after the onset of dot motion stimulation. (J-L) Geometric features of the average decision manifold: (J) Euclidean distance between corresponding arc-length points for each pair of mean paths from the two choices with equal reaction-time. (K) Cosine similarity between resolution vectors of the two mean paths, as in (J). (L) Curvature of mean paths in the contra-choice submanifold.

**Supplementary Figure 3:**
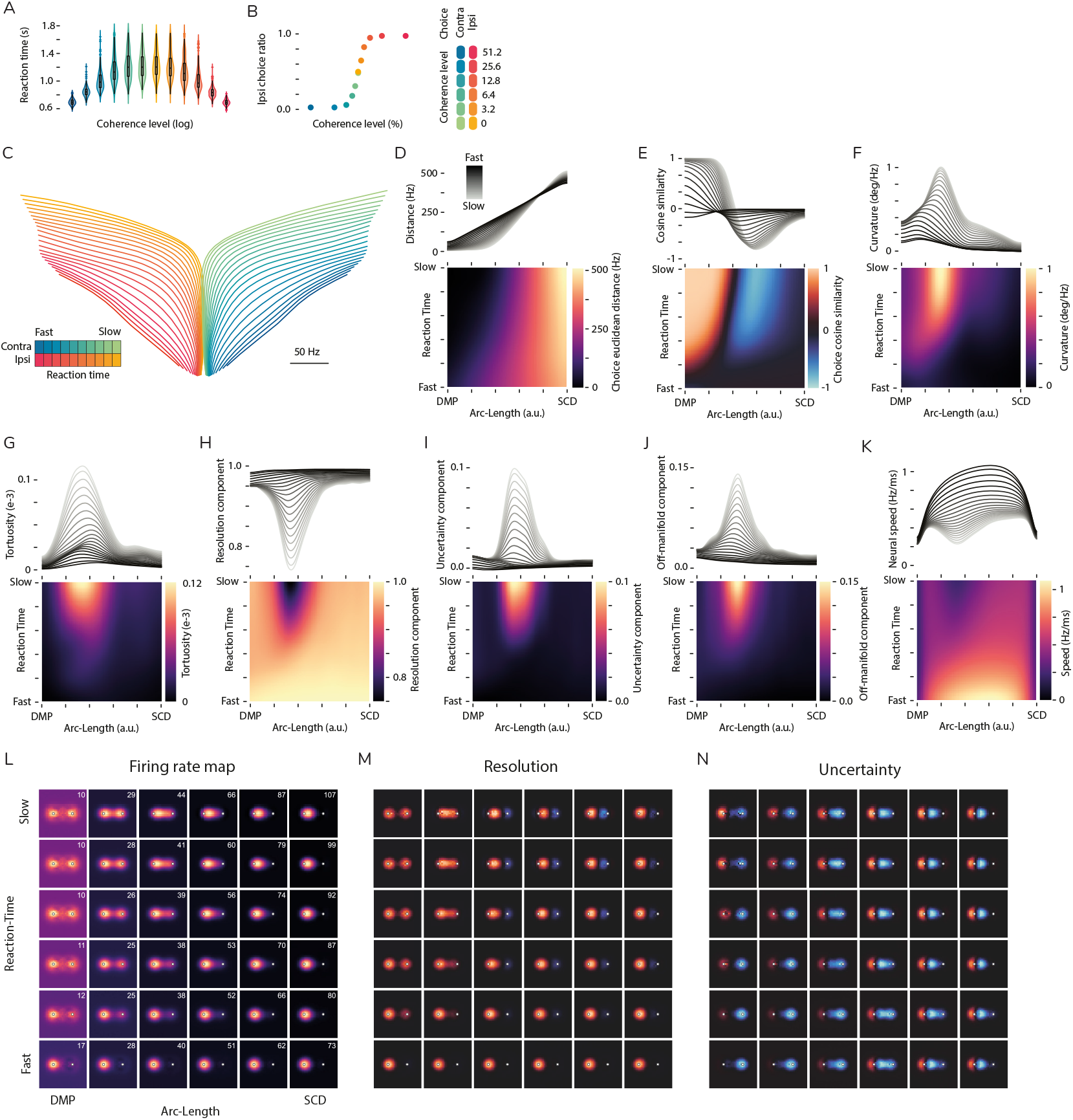
Mechanistic circuit model. **Fig. 3 Behavior and manifold geometry in the model.** (A-B) Choice selection in the model reproduces typical psychophysical curves for the distribution of reaction times (A) and the proportion of choices selecting the ipsilateral target depending on signed coherence level (B). (C) Isomap projection of the decision manifold. (D-K) Geometrical characterization of the decision manifold in high-dimensional neural (retinotopic) space. Definitions as in the main text. (L-N) Evolution of the average decision process as a function of its location on the decision submanifold for contra choices. (L) Firing rate maps illustrate how the competition between the alternatives shifts the representation across the neural sheet. Top right corner in each map displays the peak firing rate. (M-N) Resolution and uncertainty directions in retinotopic space illustrate the complex role of single neurons in driving the process forward towards resolution or integrating evidence, i.e., changing the average reaction time. The colormap is centered such that black is zero, blue represents negative values, and red represents positive values.

**Supplementary Figure 4:**
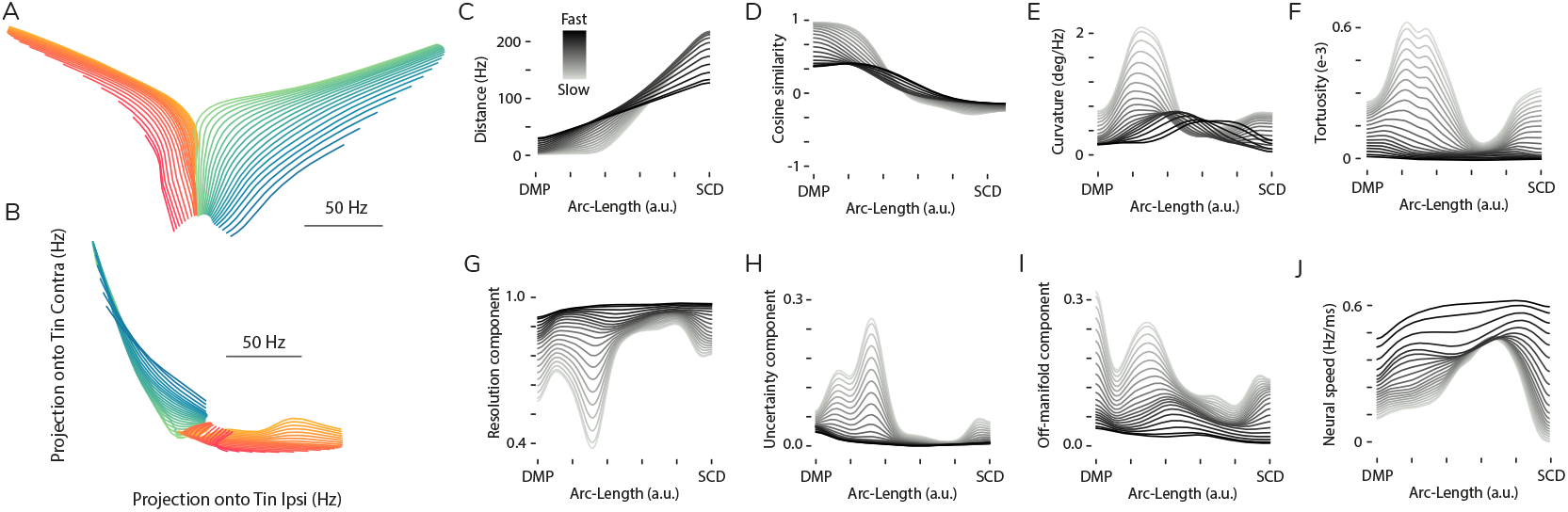
Analysis restricted to the Tin subspace. **Fig. 4 Geometry of the decision manifold projected onto the T_in_ subspace.** (A) Isomap projection of the average manifold showing robust organization by reaction time. (B) Projection of the restricted T_in_ neural activity into the plane spanned by the two vectors of neurons selective for the contra and ipsi targets. These axes can be interpreted as the classical average activity of choice selective neurons. Note the organization by reaction time largely vanishes. (C-J) Geometric characterizations of the restricted decision manifold.

**Supplementary Figure 5:**
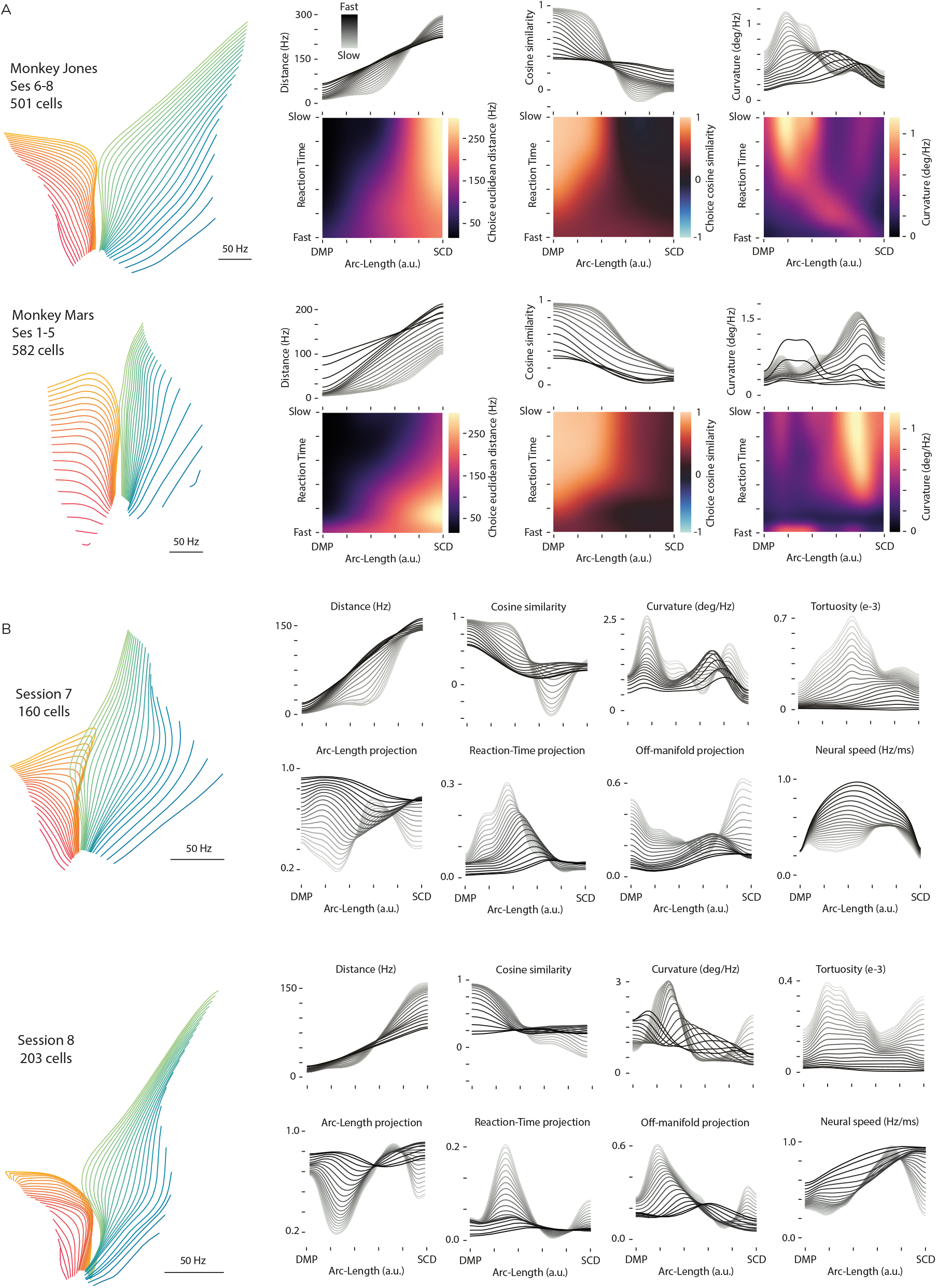
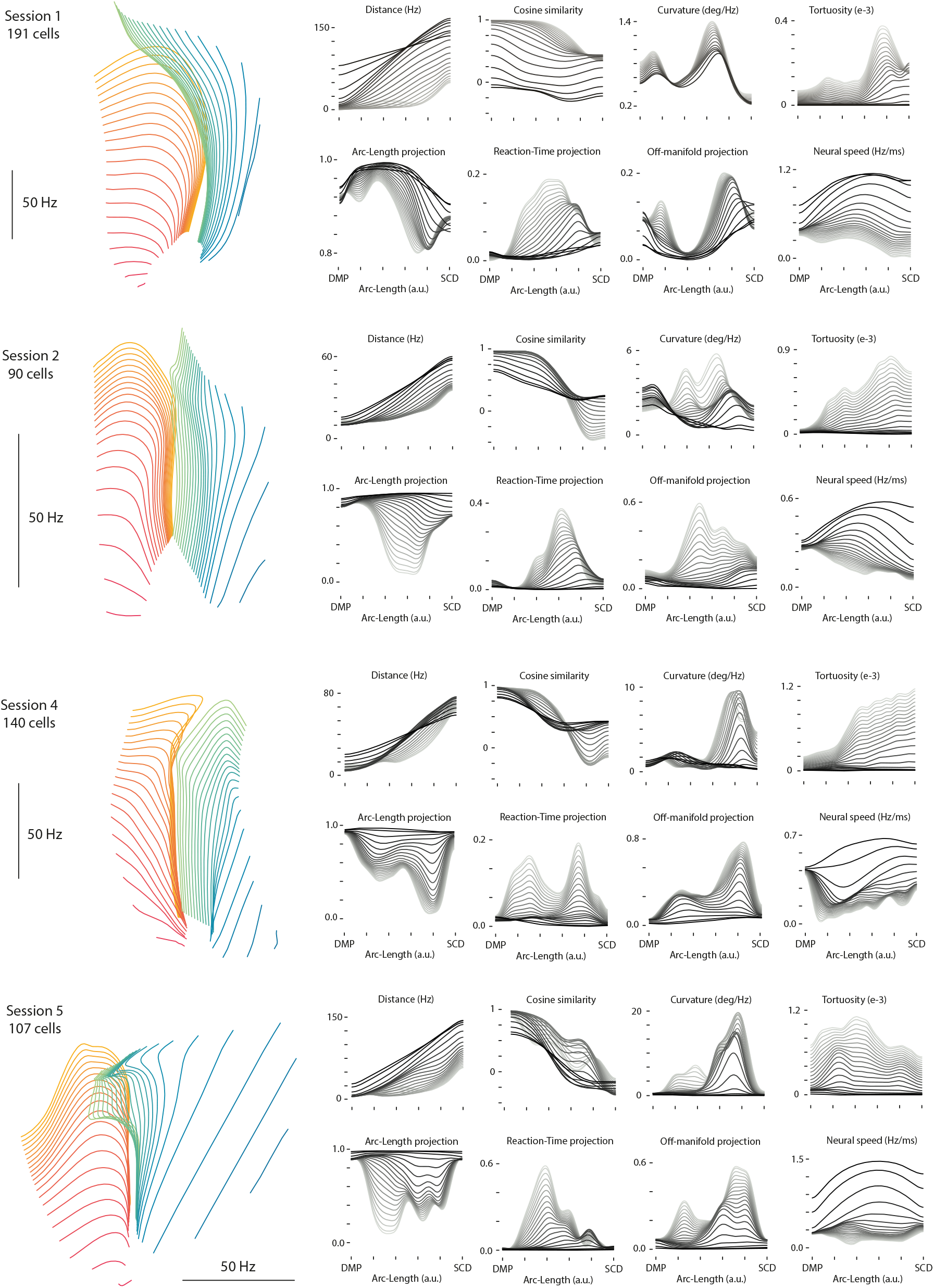
Geometry of other sessions. **Fig. 5 Geometry of decision manifolds across sessions. The example session in the main text corresponds to session 6 from Monkey Jones.** (A) Decision manifolds for each monkey. For each session, we computed the locally averaged population activity and concatenated all cells across sessions for each monkey, aligning them by arc-length and CDF-normalized reaction time. The procedure allows us to obtain an average manifold for each monkey but does not allow for single-trial analyses. (B) Decision manifold geometry for all other recording sessions, analyzed using the same approach as in the main text.

**Supplementary Figure 6:**
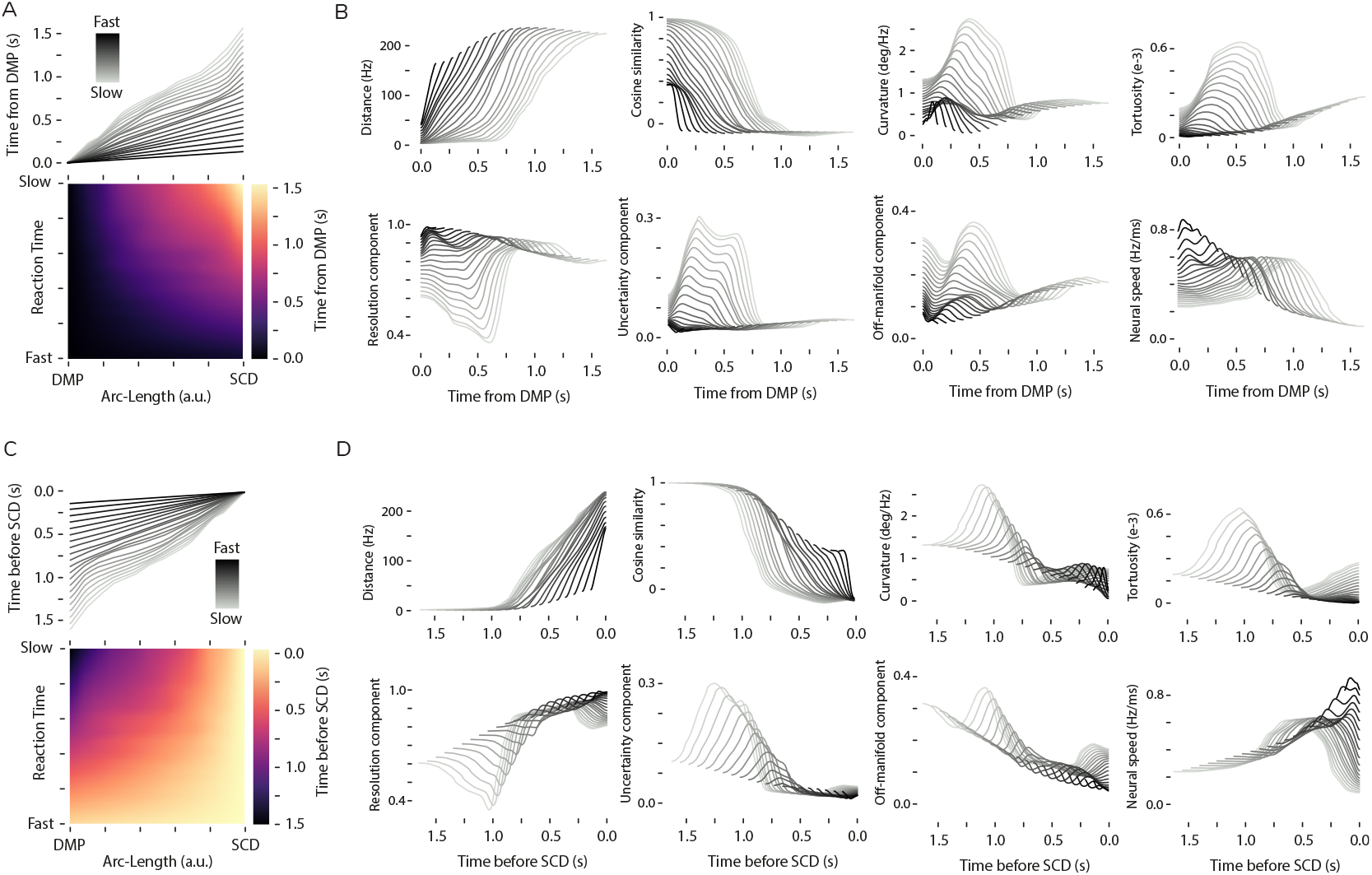
Relationship between geometry and clock-time. **Fig. 6 Geometric features in locally averaged, behaviorally aligned time.** Time is locally averaged as in Fig. 3A. (A) Locally averaged time, aligned to dot motion onset plus 200 ms (DMP). (B) Geometric features from Fig. 3 plotted against the corresponding average time from (A). (C) Locally averaged time, aligned to 60 ms before saccade initiation (SCD). (D) Geometric features from Fig. 3 plotted against the corresponding average time from (C).

**Supplementary Figure 7:**
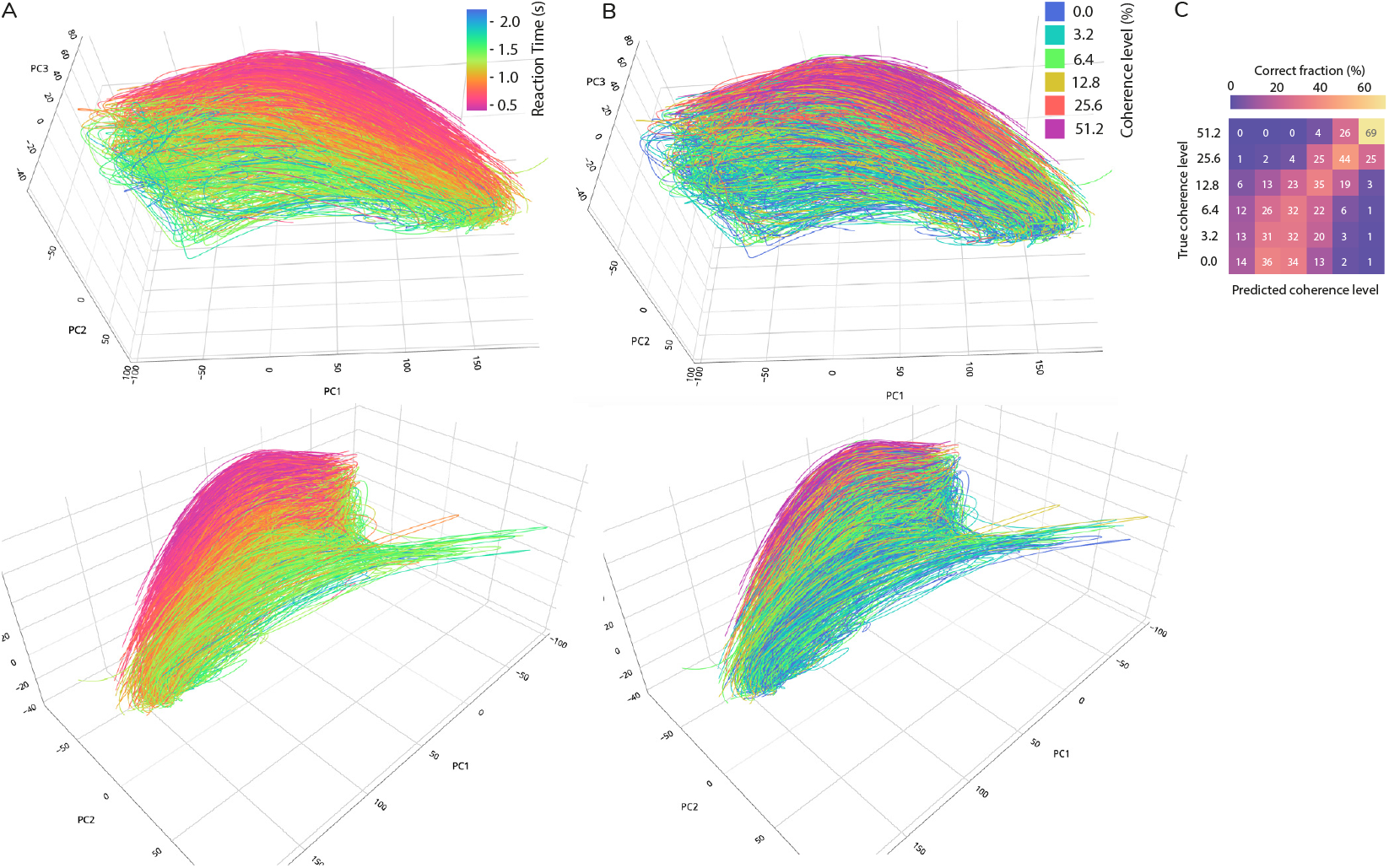
Manifold organization by coherence level. **Fig. 7 The decision manifold is less organized by motion strength.** (A) Decision manifold for the contra choice continuously colored by reaction time, analogous to the quantile-based depiction in Fig. 2A. (B) Decision manifold for the contra choice colored by coherence level, illustrating a weaker correspondence with the organization of single-trial neural state space. (C) The sequence classifier from Fig. 2C performs poorly in predicting the coherence level of individual trials based on population activity in arc-length, suggesting that coherence is less effectively captured in this state space.

